# Pedigree-tracking improved population monitoring of recessive disease-causing alleles in pedigreed canine populations

**DOI:** 10.64898/2026.10.27.761507

**Authors:** Rosalind Françoise Craddock, Mateja Janeš Fistrić, Cathryn Mellersh, Joanna Jadwiga Ilska, Pamela Wiener, Stephanie Smith, Gregor Gorjanc

## Abstract

**Background:** Most monogenic inherited diseases in dogs are recessive, so their successful control depends on genetic testing to identify carriers of the disease-causing allele. However, currently used methods for disease allele monitoring in canine populations typically only consider part of the entire pedigree, since not all dogs have a known genotype through genetic test or “hereditary status” (a determinable genotype assigned to offspring based on known parental genotypes).

**Methods:** This study uses probabilistic inference with unknown parent groups (UPGs) in five Royal Kennel Club (RKC)-registered pedigrees to estimate genotype probabilities for all individuals using available genetic test results for two monogenic recessive diseases: primary lens luxation (PLL) and progressive retinal atrophy rod-cone degeneration 4 (PRA-rcd4). From these estimates, the population trends in disease-causing allele frequencies were estimated and the inference quality was assessed through leave-one-out cross-validation and simulation.

**Results:** For dogs born after the introduction of the genetic test, the estimated disease-causing allele frequencies differed between methods. For instance, dogs born in 2020 reported up to 0.092 higher disease-causing allele frequencies with estimated genotype probabilities than those based on genetic tests and hereditary status alone. Inference performance varied across breeds, dependent on population allele frequency, with Pearson correlations between true (per genetic test result) and estimated allele dosages ranging from 0.47 to 0.67. The gene drop simulations indicated a tendency to underestimate disease-causing allele frequencies in retrospective estimates, particularly for dogs born before 1990.

**Impact:** Overall, by leveraging all available pedigree and genetic test data through probabilistic pedigree-based inference, disease-causing allele frequencies were monitored more comprehensively within the pedigreed population rather than relying on genetic test results or hereditary status alone. Such monitoring can inform responsible breeding decisions, helping to conserve genetic diversity whilst supporting improvements in future canine health and welfare.

## Introduction

Despite advances in genomics, the ongoing practice of recording parentage and maintaining pedigree records plays a critical role in the sustainable improvement and conservation of small populations [1]. With many small hobby breeders, resources often limit the application of genomic tools. Pedigree (or registered purebred) dogs provide an extensive source of pedigreed populations through the national kennel clubs [2]. While pedigree records capture genealogical relationships among dogs, genomic data provide complementary information about the genetic structure and distribution of variation within and among breeds. This combination of pedigree and genomic information is particularly relevant to understanding the distribution of inherited disease. Compared to crossbreds, pedigree dogs have higher runs of homozygosity (continuous regions of the genome that are homozygous across many loci) and are more likely to be clinically affected by a monogenic recessively inherited disease [3, 4]. The demographic history and breeding structure of modern breeds - including founder effects, genetic drift, popular sire effects, and restricted gene flow between breeds from closed registries - can result in substantial differences in genetic variation and disease-associated allele frequencies among breeds [5–10]. Consequently, many monogenic recessive diseases show breed-specific patterns of occurrence, although the demographic processes underlying these patterns may differ among diseases and breeds. For instance, primary lens luxation (PLL), a mostly autosomal recessive disease characterised by the displacement of the lens in the eye [11–14], is a recognised high risk disorder in the Miniature Bull Terrier but is rarely observed in the Standard Bull Terrier, despite the two breeds’ shared ancestry [15, 16]. The PLL-associated *ADAMTS17* allele had an estimated frequency of 27% to 39% in the Miniature Bull Terrier in 2011, whereas a recent large-scale global genomic screening study found the allele was present at very low frequency in the Standard Bull Terrier (0.32%) [4, 11, 14]. Even within breeds, some lineages will have higher disease-causing allele frequencies than others since mating is not random, thus highlighting the need for effective disease-causing allele tracking to ensure dog health, welfare, and breed conservation.

Genetic tests are available for specific hereditary diseases in dogs, many of which are caused by monogenic recessive variants. These tests target disease-associated genetic variants, with results generally presented as “clear” (no copies of the disease-associated allele), “carrier” (one copy of the disease-associated allele), or “affected” (two copies of the disease-associated allele) [17, 18]. Throughout this study, these classifications refer to the genotype at the tested locus rather than to confirmed clinical disease status. In general, the Royal Kennel Club (RKC) prioritises genetic tests targeting established causative mutations, referred to hereafter as disease-causing alleles. Genetic test results are recorded alongside the dog’s pedigree both to inform mating choices and monitor disease-associated allele frequencies over time. However, the uptake of genetic testing varies across breeds and specific hereditary diseases [17, 19]. Using available genetic tests and “hereditary status” (the propagation of genotypes to progeny from known homozygous parents), Lewis and Mellersh [17] found the frequencies of eight recessive disease-causing mutations in ten breeds were reduced by up to 90% within a decade of the genetic test introduction. However, these declines are likely overestimates for three reasons. First, less than half of RKC-registered dogs born per year have either a genetic test or hereditary status of associated locus (or loci) for at least one hereditary disease. Second, genetic testing is not random; dogs selected for breeding, including widely used sires, can contribute disproportionately to later generations [2, 20]. Hereditary status extends genotype information to additional dogs, predominantly through clear-by-clear matings that allow offspring to be assigned hereditary clear status [17]. Yet, under a fully recessive condition, clear-by-carrier matings produce no homozygous affected offspring, whilst retaining the allele in asymptomatic carriers benefiting genetic diversity. These offspring cannot be assigned a hereditary status and therefore remain unrepresented, unless genetically tested. Consequently the distribution of available genetic tests and hereditary status may not reflect the frequency of the allele across the whole registered breed population. Third, hereditary status ignores parentage error, which could introduce false-positive “hereditary clear” to dogs who are actually “carrier” or, less so, “affected”. Even a small error rate can accumulate over generations [21, 22]. Among the subset of dogs that had been genetically tested, no significant decline in mutation frequencies was observed. Thus, the decline in disease-causing allele frequency may be less when considering the whole pedigree rather than only dogs with genetic tests or hereditary status.

One solution is to estimate genotype probabilities for each individual in the pedigree using available genotype and pedigree data, also called probabilistic pedigree-based inference. Building on Elston and Stewart [23]’s application in humans, later algorithmic developments enabled accurate and computationally efficient estimation of individual genotype probabilities in large complex pedigrees, such as those in livestock [24, 25]. For example, AlphaPeel, a software for estimating (phased and unphased) genotype probabilities, achieves highly accurate estimates of individual genotype probabilities in pig, cattle, and simulated populations [26–28]. Previous studies in dogs have demonstrated early methods in a small Labrador Guide dog population (3223 individuals), including handling the pedigree loops from inbreeding [29, 30]. Although such probabilistic inference has successfully monitored allele trajectories and identified carrier status in livestock populations [31, 32], there are few published studies in smaller pedigree populations, particularly in pedigree dogs. In practice, pedigrees are sensitive to many confounding factors that influence the performance of the probabilistic pedigree-based inference. These include unknown base population allele frequencies [33, 34], the quantity and distribution of genotype data [26, 28, 35], and the pedigree completeness [27, 28]. Complete pedigrees are rare in real data. Unknown base population allele frequencies and incomplete pedigrees can be managed by grouping founders (individuals with no known parents) using “unknown parent groups” (UPG) to model possible differences among them [36, 37]. For example, Gengler *et al.* [35] grouped founders by strain and dates of birth to infer the allele trajectory of the Myostatin gene in Belgian Blue cattle achieving higher accuracy than without grouping. To improve the grouping of founders, Legarra and Aguilar [38] count the number of individuals with observed data (genotypes), called “pseudorecords”, contributing to the estimate of each UPG. Since these are additive, combining “neighbouring” UPGs can help analysts achieve a sufficient number of pseudorecords for accurate estimation of base populations parameters, such as allele frequencies.

This work studies the use of probabilistic inference with UPGs to track recessive monogenic disease-causing alleles in UK pedigree dogs. We work with five RKC-registered dog pedigrees with established causal genetic tests for autosomal monogenic recessive diseases of the eye: primary lens luxation (PLL) in the Miniature Bull Terrier, Lancashire Heeler, and Sealyham Terrier and progressive retinal atrophy rod-cone degeneration 4 (PRA-rcd4) in the Irish Setter and Gordon Setter [11, 39, 40]. These five pedigrees vary in census size, effective population size, pedigree depth, genetic test uptake, and allele frequencies for the disease variants. As these factors influence the inference quality, this allows us to evaluate the robustness of probabilistic inference in pedigrees. To this end, we (1) use state-of-the-art methods to propagate available genetic tests to all individuals in the pedigree based on the described family relationships, (2) consider different grouping strategies, pseudorecord thresholds, and genotype information (genetic test or hereditary status) for UPGs, (3) compare estimates of the disease-causing allele frequencies based on the inferred genotype probabilities with estimates based on genetic tests and hereditary status only, (4) present and discuss inference quality metrics calculated via leave-one-out cross-validation and model-based simulation, and (5) examine how the genetic test distribution affects estimates of the disease-causing allele frequency over the whole pedigree timeframe in the analysed dog pedigrees.

## Materials and methods

### Data

Information on pedigree (identifier, dam, sire, sex, and dates of birth) for five distinct breeds, along with genetic test results and hereditary status for primary lens luxation (PLL) or progressive retinal atrophy rod-cone degeneration 4 (PRA-rcd4) up to the 1st of January 2025 was provided by The Royal Kennel Club, UK (RKC) as described in Table 1. Across the five datasets, the size of pedigree ranged from 6,483 to 73,663 records, with data available as early as 1913. However, records of dogs born prior to digitisation in 1985 remain incomplete [2]. All genetic test results were obtained from commercial or academic laboratories that are recognised by the RKC using at-home saliva test kits or blood samples collected by vets, and are recorded as “clear”, “carrier”, or “affected” [41, 42]. Hereditary status is assigned to offspring of determinable genotype, which are 1) progeny of two clear parents, 2) progeny of two affected parents, or 3) progeny of one clear and one affected parent. Thus, the offspring are categorised as either “hereditary clear”, “hereditary carrier” or “hereditary affected” in the absence of a direct genetic test where the parent’s genotype associated with PLL or PRA-rcd4 is known to be homozygous (clear or affected). The available genetic test results and hereditary statuses are published for each dog by the RKC on the “Health Test Results Finder” page [43].

**Table 1.**
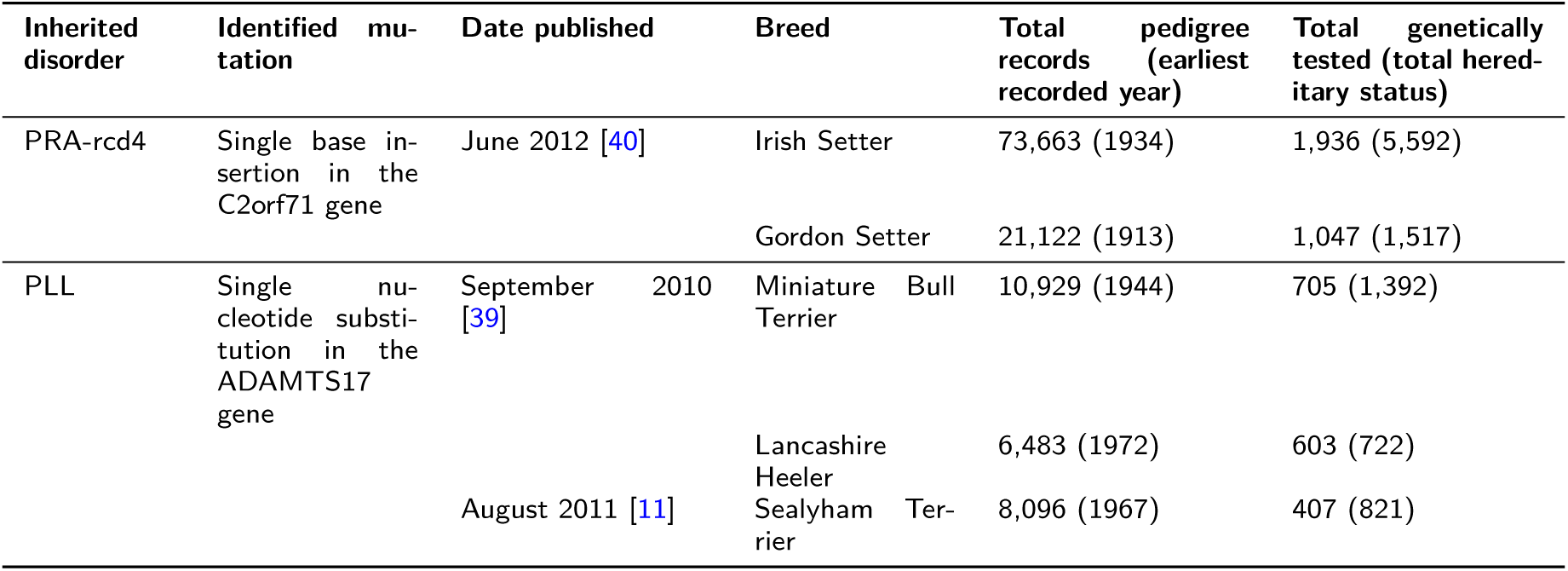
Overview of the studied breeds, disorders, and available data.

### Data cleaning

All data cleaning was completed using base R [44] and tidyverse [45]. This included the removal of any duplicates, the addition of pedigree records for any dams or sires without an individual record, and the removal of any parent records where the parent’s recorded sex was inconsistent with their designation as sire or dam. Each pedigree was sorted from ancestors to descendants. Generations were estimated forwards in time, where generation one represents the pedigree founders. For each offspring, the generation number was assigned as one greater than the highest generation number of their recorded parents. For example, a dog with parents in generation 2 and 4, generation number was calculated as max(2, 4) + 1 = 5. Thus, providing a relative measure of pedigree depth (see Supplementary Table S1). After cleaning, the genetic test missing rate ranged from 91 to 97% across the five breeds. However, this high missing rate is primarily due to the earliest genetic test publication being 2010. For dogs born from genetic test publication, the genetic test missing rate ranged from 75.9% (Lancashire Heeler) to 90.3% (Irish Setter). Figure 1 provides an overview of the proportions of the population with available genetic test results and hereditary status around the publication date of the genetic test (t) among dogs with a recorded year of birth across the five breeds, following [17].

**Figure 1.**
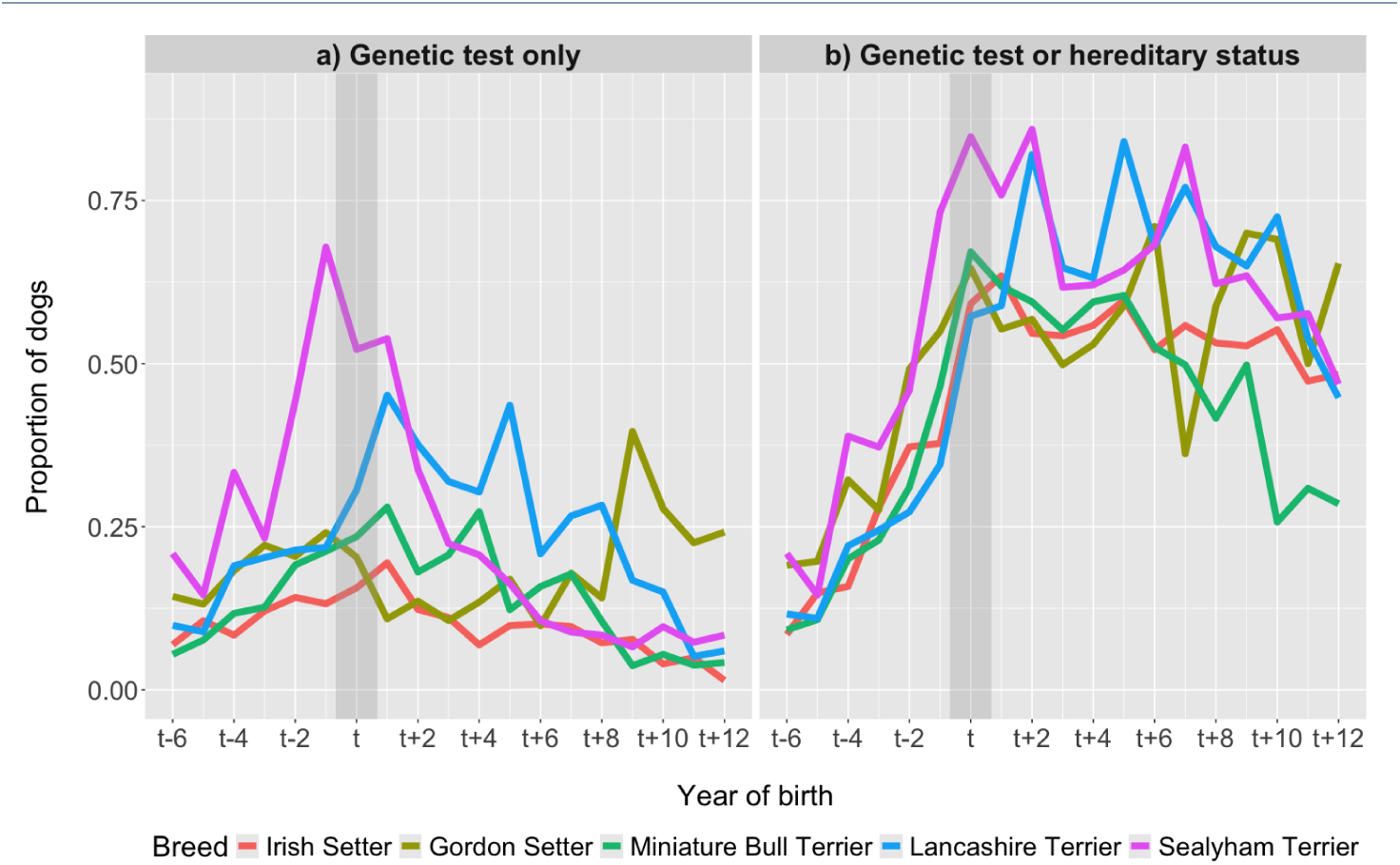
Proportion of dogs that are either a) genetically tested or b) have genetic information (genetic test or hereditary status) per year of birth for each of the five pedigrees relative to the year of the genetic test publication (t) as given in Table 1

### Assignment of unknown parent groups

All pedigrees were incomplete, with an increasing proportion of missing records back in time (see Supplementary Figure S1). Following digitisation of pedigree records, each registered imported dog was accompanied by a three-generation export pedigree, generally from an overseas kennel club recognised by the RKC, with the country of origin recorded. We used UPGs to model possible allele frequency differences in founders [36]. First, dummy individuals were created to avoid working with half-founders (individuals with only one parent known). Potential littermates were considered by assigning the same dummy individuals as parents to dogs with the same known parent and date of birth. Across the five pedigrees, the total number of unique dummy individuals created ranged from 15 (Sealyham Terrier) to 161 (Irish Setter), since half-founders represent a proportion of 0.19% (Sealyham Terrier) to 0.71% (Gordon Setter) of the whole pedigree (see Supplementary Table S1). Second, when missing, we inferred a year of birth by adding six months to the parents’ most recent birth date. Six month was used as the biological minimum generation interval. For some dogs there was insufficient information to infer year of births, thus some remained unknown: 34.1% in Sealyham Terrier, 9.87% in Irish Setter, 7.16% in Gordon Setter, 0.932% in Miniature Bull Terrier, and 0.0462% in the Lancashire Heeler. Afterwards, UPGs were assigned to each pedigree founder using one of two grouping strategies: i) birth year only and ii) birth year together with country of origin. Under the birth-year-only strategy, founders sharing the same birth year were assigned to the same UPG. Under the birth-year-and-country strategies, founders were assigned to separate UPGs for each unique combination of birth year and country of origin, resulting in a greater number of UPGs. Under both strategies, founders with an unknown birth year were included in the first UPG. The remaining UPGs were numbered sequentially according to the birth year of the founders they represented, such that earlier birth years received lower UPG numbers. Thirdly, we calculate the pseudorecords following Legarra and Aguilar [38], to count the number of genotype records that contribute to each UPG according to the pedigree. For the calculation of pseudorecords, the use of genetic test results alone was compared with genetic test results alongside hereditary status. The inclusion of hereditary status allows more UPGs per pseudorecord threshold. Finally, as the pseudorecords are additive, the “neighbouring” UPGs were combined to achieve at least 100 or 300 pseudorecords (Table 2). Thus, reducing the total number of UPGs to a maximum of 27 across the five breeds. We compared estimates of genotype and allele probabilities with or without these UPGs.

**Table 2.** Summary of the nine UPG assignments varying by grouping factor, data for pseudorecord calculation, and pseudorecord threshold for each UPG

| Grouping factor | Information used to calculate pseudorecords | Pseudorecord thresholds | Grouping strategy |
| --- | --- | --- | --- |
| Year of birth | Genetic test results only | 100 | UPG100 |
|  |  | 300 | UPG300 |
|  | Genetic test results and hereditary status | 100 | UPG100(HS) |
|  |  | 300 | UPG300(HS) |
| Year of birth and country of origin | Genetic test results only | 100 | UPGYC100 |
|  |  | 300 | UPGYC300 |
|  | Genetic test results and hereditary status | 100 | UPGYC100(HS) |
|  |  | 300 | UPGYC300(HS) |
| No UPG | n/a | n/a | No UPG |

### Probabilistic pedigree-based inference

Probabilistic pedigree-based inference was performed with AlphaPeel, a Python package to call, phase, and impute genotypes using available genotype and pedigree data [26]. This method estimates each individual’s genotype probabilities as the product of three terms: the “anterior” term is derived from parents or base population (prior) allele frequencies, the “posterior” term is informed by progeny and mates, and the “penetrance” term depends on the individual’s genotype data (genetic test result) [24–26, 46]. These terms form the Markov blanket of each individual’s genotype probabilities and are estimated iteratively by propagating (peeling) the information up and down the pedigree. To include UPGs, AlphaPeel was modified to handle new pedigree inputs and base allele frequencies for each UPG, along with adjusting the format of outputs. A new function was introduced to estimate the alternative allele frequencies for each UPG, using an expectation-maximisation approach. After each peeling cycle, these allele frequencies are estimated using the estimated genotype probabilities of the founders in each UPG (Equation 1):

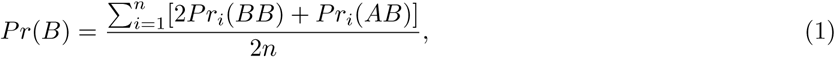

where: *Pr*(*B*) is the frequency of the disease-causing allele (B) in a cohort, *Pr_i_*(*BB*) is the estimated homozygous recessive (affected) probability for individual *i* in a cohort, *Pr_i_*(*AB*) is the estimated heterozygous (carrier) probability for individual *i* in a cohort, and *n* is the number of individuals in a cohort. This enhancement was tested for accuracy and efficiency, and implemented in AlphaPeel release version 1.3.0. For each of the five registered breeds, single-locus analysis was performed across five peeling cycles (the default) using two input files: pedigree file (including UPGs) and genotype file (containing only genetic test results). The initial allele frequencies for the “anterior” term were estimated using observed genetic data through the Newton method [e.g., 47]. Then, the allele frequencies were updated for each UPG after every peeling cycle. The “penetrance” term was modelled with a single genotype error rate, updated after each peeling cycle based on the mean probability that the estimated genotype of tested individuals does not match the observed results, to absorb some Mendelian inconsistencies such as from sampling errors, laboratory errors, mutations, or, less so, pedigree error. Erroneous parent-offspring relationship, although likely present, is not directly modelled. Therefore pedigree errors may still propagate information through the wrong pedigree link, unless a Mendelian inconsistency occurs.

### Estimation of disease-causing allele frequencies

The estimated individual genotype probabilities were averaged by birth year, and the population allele frequency trends were calculated using Equation 1 per birth year. For comparison, we also estimated the disease-causing allele trajectories using the available genetic data and hereditary status using the naïve estimator as in Lewis and Mellersh [17]:

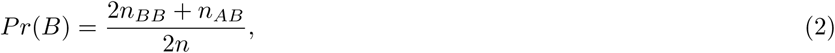

where: *Pr*(*B*) is the frequency of the disease-causing allele (B) in a cohort, *n_BB_* is the number of individuals recorded as homozygous for the disease-causing allele (affected) in the cohort, *n_AB_* is the number of individuals recorded as heterozygous (carrier) in the cohort, and *n* is the number of individuals with genetic data and hereditary status data in the cohort.

The 95% confidence intervals for both estimates were obtained by bootstrapping with 2,000 replicates. In each replicate, dogs were sampled with replacement within each year of birth from 2000 onwards. For genotype probabilities, sampling was performed from all dogs born in each year, whereas for genetic test results and hereditary status, sampling was restricted to dogs born in each year with available genotype information.

### Quality of estimates

Two different approaches were used for assessing the quality of the estimates: leave-one-out cross-validation and simulation.

### Leave-one-out cross-validation

All the observed genetic test results were masked from the inference one at a time. Then, comparisons were made between the true (observed, but masked) genetic test results and the estimated genotype probabilities or allele dosages. As only one genetic test result was removed each time, this minimised the impact of missing genotypes on the results [28, 48]. Various metrics were used to assess the overall quality of estimation. To measure accuracy, the Pearson correlation was used between the true allele dosage and the estimated allele dosage when masked. To measure reliability, the imputation quality score (IQS) was used [49, 50]. Table 3 describes the multi-categorical confusion matrix used by IQS through the Equations 3, 4, & 5.

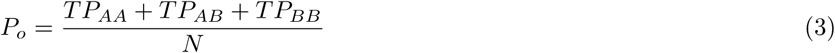

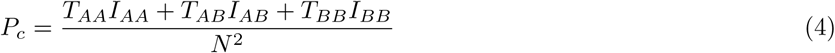

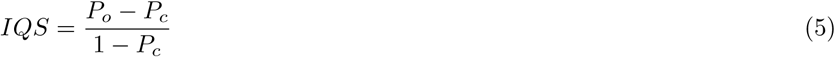

**Table 3.** Multi-class genotype concordance for the three genotype states AA (“Clear”), AB (“Carrier”), and BB (“Affected”) where: *TP_X_* is the true positive metric calculated as the sum of *X* genotype probabilities across individuals with *X* genetic test result, *T_X_* is the number of true genotypes *X*, *I_X_* is the number (or sum of probabilities) of inferred genotypes *X*, and *N* is the total number of individuals with a genetic test

|  | True genotype |  |  | Total inferred |
| --- | --- | --- | --- | --- |
|  | AA | AB | BB |  |
| Inferred AA | $TP_{AA}$ | | | $I_{AA}$ |
| Inferred AB | | $TP_{AB}$ | | $I_{AB}$ |
| Inferred BB | | | $TP_{BB}$ | $I_{BB}$ |
| Total true | $T_{AA}$ | $T_{AB}$ | $T_{BB}$ | $N$ |

By separating the concordance rate, *P_o_*, and the chance agreement, *P_c_*, IQS accounts for allele frequency. Hence, providing a robust comparison amongst the five breeds. Alternative reliability metrics, including squared Pearson correlation, were calculated but were not presented. To assess the performance of estimates by genotype state, the calibrated F1 score was calculated from the binary confusion matrix of each genotype state: clear (AA), carrier (AB), affected (BB), using the probabilistic counts as described in Table 4 and Equations 6, 7, 8, 9, & 10 [51, 52].

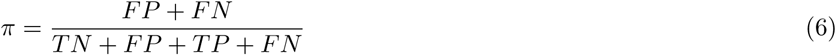

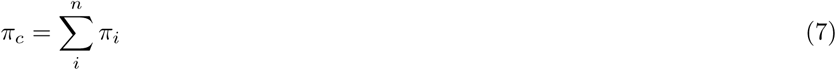

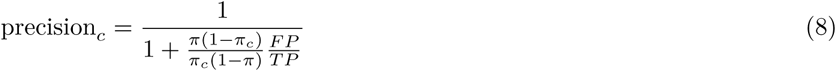

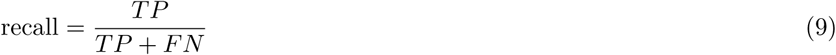

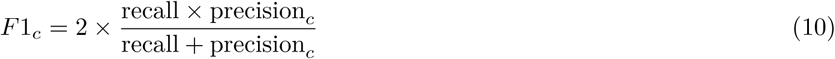

**Table 4.** Binary confusion matrix used for calculation of calibrated precision (Equations 6, 7, & 8), recall (Equation 9), and calibrated F1 score (Equation 10)

| Inferred class | True class |  |
| --- | --- | --- |
|  | Positive class (e.g AB or BB) | Negative class (e.g not AB or not BB) |
| Predicted positive<br>(e.g AB or BB) | True positive (TP) | False positive (FP) |
| Predicted negative<br>(e.g not AB or not BB) | False negative (FN) | True negative (TN) |

The F1 score was calibrated by *π_c_*, which represents the average proportion of positive class (carrier or affected) among the available genetic test results across the five breeds (Equations 6 & 7; Siblini *et al.* [52]). By calibrating, the F1 score accounts for class imbalance, common with genetic disorders. For example, there are far fewer genetic test results of affected individuals compared to either clear or carrier. Thus, ensuring fair comparison across breeds and genotype state.

To assess systematic bias in estimated allele dosages, the mean signed dosage error was calculated separately for each true genotype state. For each dog with a genetic test result, the estimated allele dosage was subtracted from the masked true genotype to obtain the signed error. The mean sign error was then calculated within each true genotype state. Positive values indicate overestimation of allele dosage, whereas negative values indicate underestimation [53]. 95% confidence intervals for the mean signed error were calculated per Equation 11.

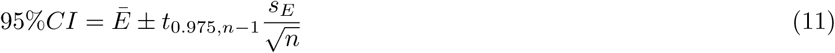

where: *Ē* is the mean signed error, *s_E_* is the standard deviations of the errors, *n* is the number of tested dogs with the corresponding true genotype state, *t*_0.975*,n*_*_−_*_1_ is the two-sided t critical value with *n −* 1 degrees of freedom.

### Model-based simulation

We used forward-in-time (gene drop) simulations [54] to evaluate the quality of estimates across the entire pedigree time-frame. First, all founders were assigned a discrete genotype using their individual estimated genotype probabilities from the UPGYC100 model (founders grouped by year of birth, country of origin, and a pseudorecord of 100 or more; Table 2). Thus, prioritising internal consistency over independent validation for the founders whilst allowing realistic simulation of empirical allele frequencies and founder diversity. Next, these assigned alleles were dropped to offspring using the available pedigree and Mendelian inheritance without mutation. As we expect breeders to select against the disease-causing allele, selection was applied to dogs with pre-existing genetic test or hereditary status recorded before the birth of their earliest offspring. For these dogs, the favourable genotype (the genotype free of the disease-causing allele) was preferentially assigned from the possible parental mating outcomes, rather than through random assignment. For simplification, the gestation length was omitted from the criterion, as offspring birthdate was used in place of the conception date; a difference of approximately 63 days [55, 56]. A simplification which was deemed sufficient for the study’s purpose. The simulation proceeded through each subsequent generation to generate genotypes for all individuals in the pedigree. We repeated the simulation 100 times giving us different genotype realisations for the real pedigree and estimated genotype probabilities of pedigree founders.

To understand the influence of genetic test distribution and possible bias of the estimated disease-causing allele frequencies, we evaluated three different scenarios. These three scenarios, “Observed”, “Even”, and “Bias”, were designed to differ only in the distribution of observed genotypes within each pedigree, while maintaining the target number of observed genotypes as consistent as possible across scenarios. For example, the Gordon Setter has 1047 genetic tests for PRA-rcd4 in the real data and was therefore assigned a target of 1047 observed genotypes in each scenario. These observed genotypes were obtained independently from each of the 100 simulations in line with the total number of existing genetic test results for the corresponding disease and breed. However, the target number could not always be achieved exactly because of constraints imposed by the sampling procedure. Consequently, the actual number of observed genotypes varied slightly among scenarios and simulation replicates, as described below.

The first scenario (“Observed”) distributed the genotypes to match the distribution of genetic tests in the real data. Although this holds a genotype for the same individuals as the real data, the genotype can vary amongst the simulated repeats due to randomness of gene dropping. Thus, this scenario retained the empirical distribution and sample size of genetic testing while allowing the underlying genotypes to vary among simulations.

The second scenario (“Even”) distributed the target number of observed genotypes across generations in proportion to the number of animals in each generation, resulting in approximately equal proportion of animals being genotyped in each generation. Within each generation, the distribution of observed genotypes was random. Thus, it did not consider families within generations. Where the target number of observed genotypes could not be divided evenly across the generations, the same number of observed genotypes was used across all simulation replicates, but was slightly lower than the target. In all breeds, the resulting number of observed genotypes were at least 95% of the target number.

The third scenario (“Bias”) considered sampling bias (high number of affected dogs to identify the disease-causing allele) and ascertainment bias (preferential recording of clear dogs after genetic test introduction). Although both biases were considered plausible in the available data for certain conditions and/or breeds [17], the scenario represented an extreme manifestation of these potential biases. Initially, individuals were separated into three groups: before testing, prior to genetic test introduction, and after genetic test introduction. The before testing groups comprised of all dogs born before the year of the earliest born genetically tested dog. The prior to genetic test introduction holds all dogs born after the year of the earliest born genetically tested dog up until genetic test publication per Table 1. The after genetic test introduction group contains all dogs born from the year of test publication onwards. Next, we checked the split of the genetic tests collected prior to and after genetic test introduction to obtain the total number of observed genotypes in each group. Afterwards, we set 60%, 20%, and 20% of observed genotypes prior to genetic test introduction respectively as affected, carrier, and clear. After the genetic test introduction, we set 80% clear, 15% carrier, and 5% affected. This proportion was enforced, sacrificing the total number of observed genotypes in the pedigree in some simulated repeats with insufficient number of clear, carrier, or affected genotypes. Consequently the sample size of observed genotypes varied among replicates. Relative to each breed’s target, the range in replicates was 61-109% in the Irish Setter, 53-116% in the Gordon setter, 88-101% in the Miniature Bull Terrier, 9.45-114% in the Lancashire Heeler, and 15-101% in the Sealyham Terrier. Through all breeds, the replicate median was 87% of above. The chosen proportions represent reasonable extremes, however, it is not representative of exact patterns since these are unknown. Thus, provided a challenging sampling configuration under which to assess the sensitivity of estimated disease-causing allele frequencies to non-random genetic test distributions.

Following the preparation of observed genotype files for the three scenarios, AlphaPeel was used to calculate genotype probabilities using the real pedigree with UPGYC100 and to estimate allele frequencies by year of birth. We then calculated the difference between the true disease-causing allele frequency from the simulations and the estimate from AlphaPeel for each of the 100 simulation replicates. We hypothesised that the different distribution of genotypes in the three scenarios causes differences in the estimated disease-causing allele frequencies over time. We tested this for each pedigree with the mixed effects model in Equation 12, fitted using the lmer R package [57] separately to each year born:

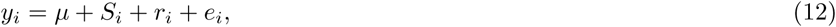

where: *y_i_* is the allele frequency difference (true – estimated) for the *i*-th replicate, *µ* is the intercept representing the “Observed” scenario, *S_i_* is the scenario factor for the “Even” and “Bias” scenarios, *r_i_* is the random deviation of simulation replicate accounting for shared simulated genotypes from shared pedigree data, assuming 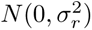, and *e_i_* is the random residual, assuming 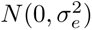. Due to large number of statistical tests, p-value thresholds for each pedigree were adjusted using the Bonferroni method, and differences were assume significant when a p-value was equal to or smaller than 0.05*/n*, where *n* is the number of years in the pedigree.

All scripts used for this study are available through the GitHub repository: https://github.com/HighlanderLab/RCraddock_AlleleFetch_Geno.

## Results

### Comparisons of allele trajectories across UPG assignments

Figure 2 compares the impact of different UPG assignments on the estimated allele frequencies by year of birth for the founders (A) and all dogs in the whole pedigree (B) for PLL in the Miniature Bull Terrier (column 1) and for PRA-rcd4 in the Gordon Setter (column 2). Although we evaluated eight different UPG assignments based on year of birth, country of origin, and number of pseudorecords using genetic tests only or also hereditary status (Table 2), we only present the four that provided consistently different results: UPG100, UPGYC100, UPGYC300, and UPGYC100(HS).

**Figure 2.**
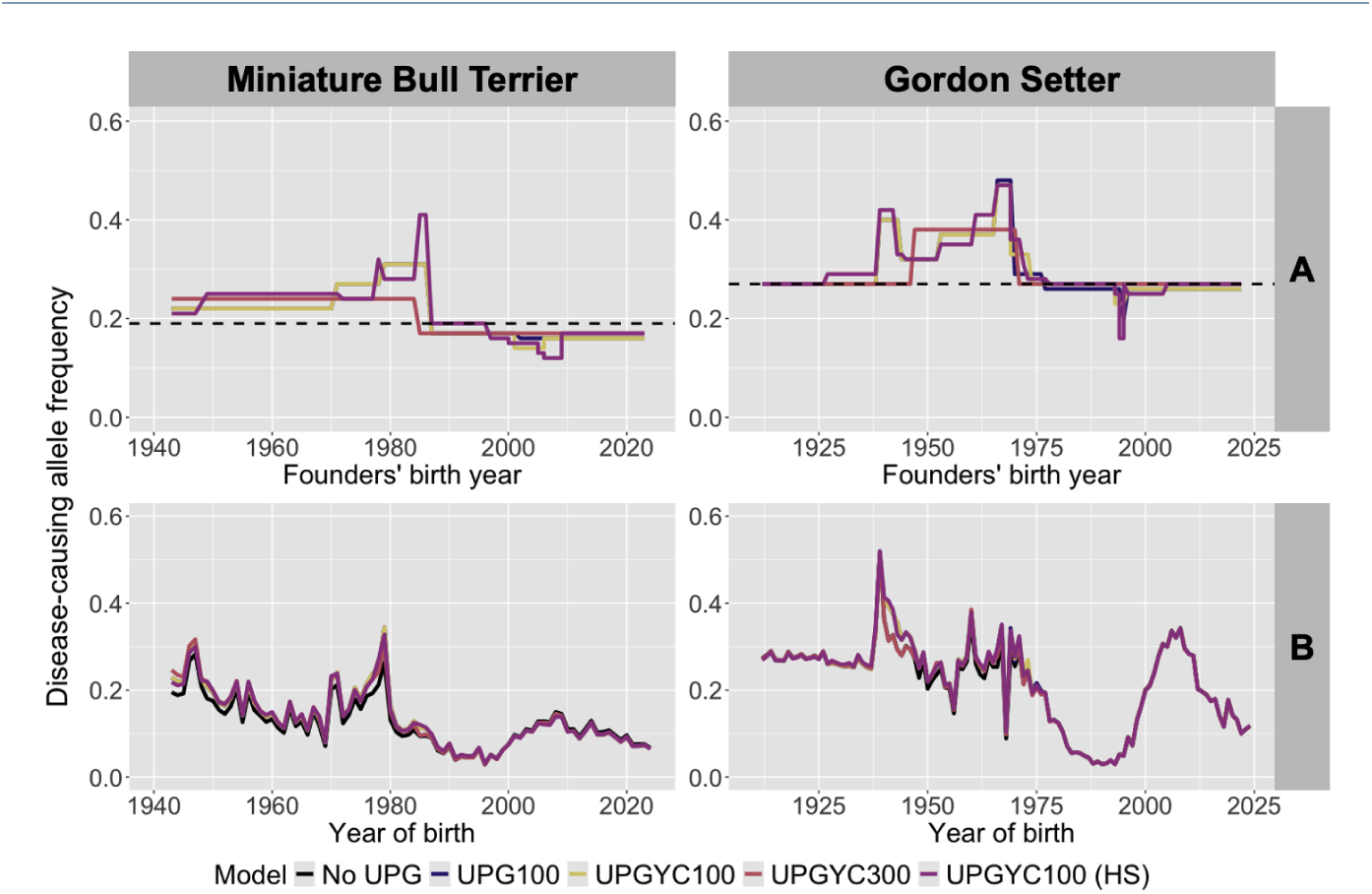
Estimated allele frequencies with different UPG assignments A) across the founder’s birth years and B) across the whole pedigree birth years without UPG, UPG100, UPGYC100, UPGYC300, and UPGYC100(HS) in the Miniature Bull Terrier (column 1) and Gordon Setter (column 2).

UPG100 (blue line) gave only 5 UPGs in the Miniature Bull Terrier and 9 in the Gordon Setter. The UPG100 trend (blue line) was often obscured by the UPGYC100 (gold line) on the graph, reflecting that adding country of origin made little or no difference to the allele frequencies across the founders (A) and the entire pedigree (B). In the Miniature Bull Terrier, UPGYC100 provided an additional UPG separating Australia from Europe in 2001 compared to UPG100. UPGYC100 indicated that the frequency of the PLL-causing allele was lower in imported Australian-born Miniature Bull Terriers than in European-born Miniature Bull Terriers in 2001 (Figure 2A, gold line). The Gordon Setter had the same number of UPGs whether UPG100 or UPGYC100 assignment was used. However, the assignment differs in founders’ born from 1960 to 1990 (Figure 2, blue and gold lines). For example, all founders born in 1969 were assigned UPG 5 with UPG100 but for UPGYC100, two dogs originating from Norway were separated into UPG 6. Across all the pedigrees, the number of UPG assigned with UPG100 or UPGYC100 ranged from 3 (Sealyham Terrier) to 12 (Irish Setter).

UPGYC300 had the least number of UPGs assigned, ranging from 1, equivalent to no UPG (Sealyham Terrier and Lancashire Heeler), to 5 (Irish Setter). In the Miniature Bull Terrier and the Gordon Setter, UPGYC300 respectively gave 2 and 3 UPGs (Figure 2, pink line). These UPGs captured allele frequency differences across founders; for example 0.05 difference in the Miniature Bull Terrier. UPGYC100(HS) provided the highest number of UPGs, ranging from 6 (Lancashire Heeler) to 27 (Irish Setter). In the Miniature Bull Terrier and the Gordon Setter using UPGYC100(HS) captured a range of 0.3 in allele frequencies across UPGs, with 14 and 15 UPGs. With more UPGs, there are fewer founders in each grouping. This may explain some of the extreme changes observed with UPGYC100(HS), such as between 1983 and 1987 in the Miniature Bull Terrier (Figure 2).

Evaluating allele frequency trends across the whole pedigree, most of the difference observed with UPGs compared to no UPG (black line) was observed in the early years of the pedigree (Figure 2B). This was as expected since most of the founders were born early in the pedigree, and these are the furthest from the available genotype data with the highest uncertainty, as shown in Supplementary Figure S2 using model UPGYC100. Generally prior to 1990 (the early pedigree), the use of UPG increased the estimates of disease-causing allele frequency. In more recent years, there was either no difference with UPG assignment compared to without (Gordon Setter, Sealyham Terrier, and Lancashire Heeler) or using UPGs slightly reduced estimated disease-causing allele frequencies after genetic test introduction (Miniature Bull Terrier and Irish Setter, results not shown).

Although we observe only subtle differences in estimated allele frequency trends with and without UPGs, the UPGs enables the estimation of different allele frequencies across time and country of origin amongst the founders. How the UPG is assigned can lead to differences in the allele frequencies represented by each UPG, but has smaller influence on the allele trajectories across the whole pedigree. For the remainder of the paper, we present results using UPGYC100 assignments.

### Comparisons of estimated disease-causing allele frequencies

Using the UPGYC100, the disease-causing allele frequency trends were estimated from genotype probabilities across the whole pedigree and compared to estimates using the portion of the pedigree with a genetic test result or hereditary status. Figure 3 compares these two estimates for each of the five breeds with 95% confidence intervals obtained via bootstrapping. In dogs born in the early 2000s, before the publication of the genetic tests, the inferred genotype probabilities estimated, on average, a lower disease-causing allele frequency than with genetic test results and hereditary status. The high fluctuations in estimates based on genetic test and hereditary status and wide confidence intervals reflect the small sample size of individuals with genetic test or hereditary status per birth year. In the Miniature Bull Terrier and Sealyham Terrier, this is as low as 1 in the year 2000. With the estimated genotype probabilities, all individuals in the pedigree contribute probabilistically to the disease-causing allele frequency trends, rather than restricting the analysis to individuals with available genetic test results or hereditary status. This provides greater pedigree-based population coverage with a minimum of 45 individuals per birth year in the Sealyham Terrier. Although the Sealyham Terrier is not the smallest-sized pedigree of the five breeds, they have a high rate of missing dates of birth (43%), with only some inferrable from the data. Thus, the trends by birth year represent around 66% of the Sealyham Terrier pedigree. This was due to a high number of imported dogs with either no recorded descendants or no descendants with recorded dates of birth. Supplementary Figure S4 presents the estimated allele frequency trends from genotype probabilities by generation, thus captures all recorded individuals in the pedigree information for each breed.

**Figure 3.**
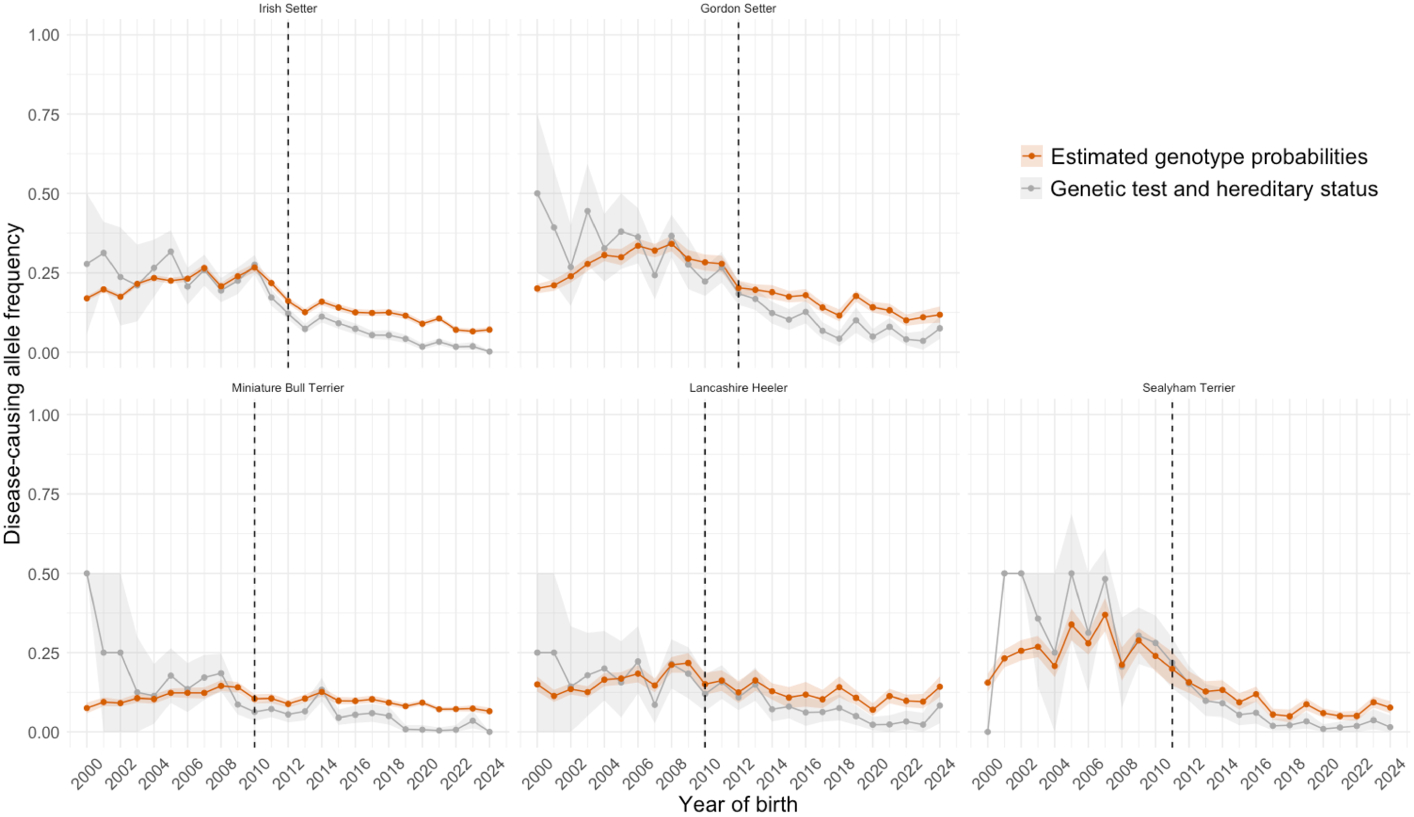
Comparison of estimated disease-causing alleles by year of birth using either the genotype probabilities in orange or available genetic test results and hereditary status in grey for PLL in Miniature Bull Terrier, PRA-rcd4 in Irish Setter, PLL in Lancashire Heeler, PRA-rcd4 in Gordon Setter, and PLL in Sealyham Terrier. The ribbons capture the 95% confidence interval obtained from bootstrapping.

In nearly all years after the introduction of genetic tests, using genotype probabilities gave a higher estimate of disease-causing allele frequency than genetic tests and hereditary status for all five breeds. This difference is up to 0.092 in the Gordon Setter in 2020. Although the difference is smaller in the other breeds, this trend is consistent across all five breeds.

### Estimation of genotype frequency trends

In addition to the allele trajectories, the estimated genotype probabilities enabled us to track temporal changes in unphased and phased genotype frequencies beyond what is possible from the observed genetic test results and hereditary status alone. Figure 4 present the genotype frequencies from birth years 2000 to 2025 using the estimated genotype probabilities for the respective disease in all five pedigrees. The “clear”, “carrier”, and “affected” represents the unphased genotype frequencies, while the “maternal carrier” and “paternal carrier” represents the phased genotype frequencies of heterozygotes. In all five pedigrees, the frequency of the clear genotype increased from around the genetic test introduction, reducing the frequency of the carrier and affected genotypes. Prior to genetic test introduction, most disease-causing alleles in the carriers (heterozygotes) were inherited from the sire (paternal carrier), whereas afterwards this switched to the dams (maternal carrier). Such fine-scale population trends can signal whether the estimated disease-causing allele persists in the population primary among carrier dogs or also present in dogs with an affected genotype. For example, although the Gordon Setter had a 0.12 disease-causing allele frequency for PRA-rcd4 in the 2024 birth cohort (Figure 3), nearly all were present in carriers (heterozygotes) and not affected genotypes according to our estimates (Figure 4). Thus, assuming complete recessivity and disregarding potential genetic heterogeneity, very few dogs born in 2024 would be expected to develop PRA-rcd4 at around 10 years of age. As described in the first additional file of the supplementary materials and shown in Supplementary Figure S3, the estimated genotype frequency trajectories can be plotted across the whole pedigree time-frame with 95% approximate credible intervals to capture uncertainty too.

**Figure 4.**
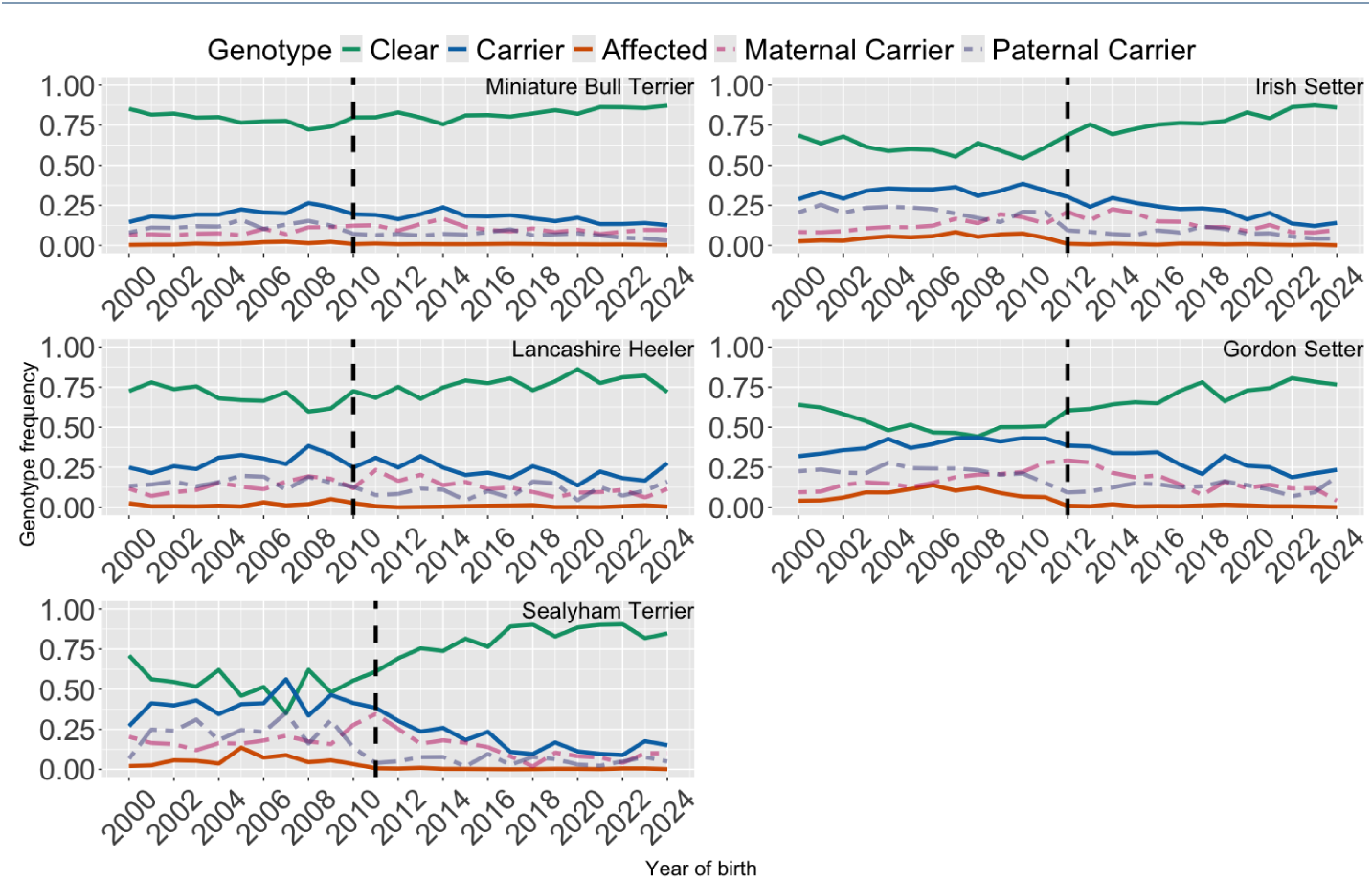
Estimated phased genotype frequencies by year of birth from 2000 using the genotype probabilities for PLL in Miniature Bull Terrier, PRA-rcd4 in Irish Setter, PLL in Lancashire Heeler, PRA-rcd4 in Gordon Setter, and PLL in Sealyham Terrier.

### Estimation quality through leave-one-out cross-validation

The overall inference quality is presented through accuracy (Pearson correlation) and reliability (IQS) in Table 5. Despite the different models of UPG assignment tested, no difference was observed between them. Hence, we only present results for the UPGYC100 assignment. Using a leave-one-out cross-validation, we compared the estimated allele dosage or genotype probabilities, when one genotype was masked, to the true genotype as defined by the genetic test result. Moderate to high quality of inference was achieved across all five breeds. The difference between the highest accuracy (Gordon Setter) and the lowest (Miniature Bull Terrier) was 0.21. When ranking breeds, the Pearson correlation matched the order of population disease-causing allele frequencies in the breeds. With accounting for these disease-causing allele frequencies, the IQS reports a different ranking, with the Sealyham Terrier having a lower reliability than the Lancashire Heeler and similar to the Miniature Bull Terrier. Both Sealyham Terrier and Miniature Bull Terrier have relatively higher pedigree effective population sizes [6]. The Sealyham Terrier had imports of up to 40%, whereas the Miniature Bull Terrier had relative low proportion of known ancestors up to five ancestral generation around genetic test introduction (see Supplementary Figure S1). Both factors may contribute to lower reliability (by IQS) in estimated genotype probabilities.

**Table 5.** Inference accuracy (Pearson correlation) and reliability (imputation quality score) for the five breeds with disease-causing allele frequency (ordered from highest to lowest accuracy).

| Breed | Pearson correlation | Imputation quality score | Disease-causing allele frequency |
| --- | --- | --- | --- |
| Gordon Setter | 0.67 | 0.36 | 0.27 |
| Irish Setter | 0.59 | 0.28 | 0.25 |
| Sealyham | 0.53 | 0.25 | 0.25 |
| Lancashire Heeler | 0.49 | 0.28 | 0.21 |
| Miniature Bull Terrier | 0.47 | 0.25 | 0.20 |

Inference quality was not uniform across genotype states. Across all five breeds, the highest performance, as measured by the calibrated F1-score, was observed for the clear genotype state (Table 6). In contrast, performance for carrier genotype was moderate, while performance for affected genotypes were moderate to low. It is easier to identify clear dogs than carrier or affected dogs, as the clear genotype state exists in higher frequencies in the population. Given the similarity of the F1-score for the clear genotype class across the five breeds, the genotype-specific results indicate that differences in overall inference quality among breeds were driven less by the ability to identify clear dogs and more by the ability to correctly infer the affected genotype. In particular, the Lancashire Heeler, had the lowest calibrated F1-score for the affected genotype, nearly 0.1 lower than the next lowest-performing breed, the Sealyham Terrier. However, the number of dogs genetically tested as affected was low in the Sealyham Terrier, Lancashire Heeler, and Miniature Bull Terrier, with only 9 or 10 (Table 6). Consequently, performance estimates for the affected genotype in these breeds were less precise.

**Table 6.**
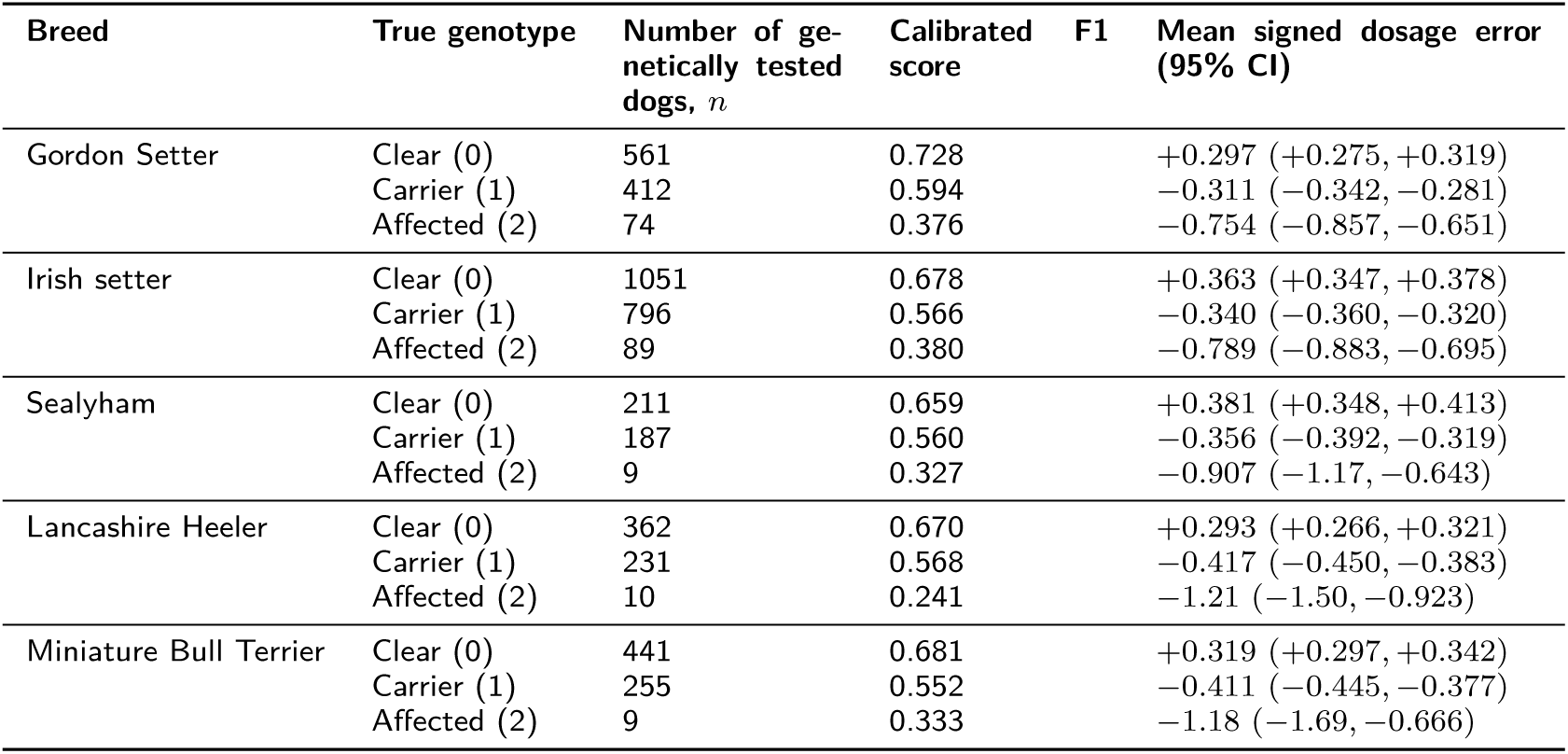
Genotype-specific calibrated F1-score and mean signed dosage error for each breed. The F1-score was calibrated by the mean genotype state proportions across all five breeds.

The mean signed dosage errors (Table 6) indicated a general compression of estimated allele dosages towards intermediate values. Dogs with true clear genotypes were marginally overestimated, whereas dogs with true carrier or affected genotypes were underestimated. Some movement away from the extreme dosage values of 0 and 2 was expected due to the inclusion of uncertainty through the genotype probabilities. However, the magnitude, particularly for the dogs with a true affected genotype, indicates a degree of systematic shrinkage. Based on the genotype-specific errors and the number of dogs genetically tested in each genotype class, allele frequencies were estimated to be marginally overestimated in the Irish Setter and Miniature Bull Terrier (Supplementary Table S2). As the 95% confidence intervals of the Gordon Setter, Sealyham Terrier, and Lancashire Heeler contained zero, there was no clear evidence of systematic over- or underestimation in the genetically tested cohort of these breeds.

### Model-based quality assessment via simulations

Figures 5 and 6 present the difference between the true and estimated allele trajectories for “Observed”, “Even”, and “Bias” distributions of genotypes using simulated data, each with 100 replicates. With the genotypes distributed per reality (“Observed”), there was a tendency to underestimate the disease-causing allele frequency in all pedigrees (true *>* estimated allele frequency). Initially, there is high uncertainty in the estimates, which decreases over time as genotype information is incorporated, as shown by the reduction in variation among the 100 replicates. Agreeing with findings presented in Supplementary Figure S2. As expected, when the genotypes were distributed consistently within all generations of the pedigree (“Even”), the difference between estimate and true were centered around zero with decreasing variance over birth years as genotypes accumulated with each successive generation. When considering the influence of sampling for affected dogs followed by preferential reporting of clear dogs (“Bias”), we observed a larger variance amongst the simulated replicates, but no consistent agreement, overestimation, or under-estimation of allele frequencies was present across all five breeds. This highlights the differences in allele frequency estimation using genotype probabilities from the distribution of genetic tests within the various pedigree structures.

**Figure 5.**
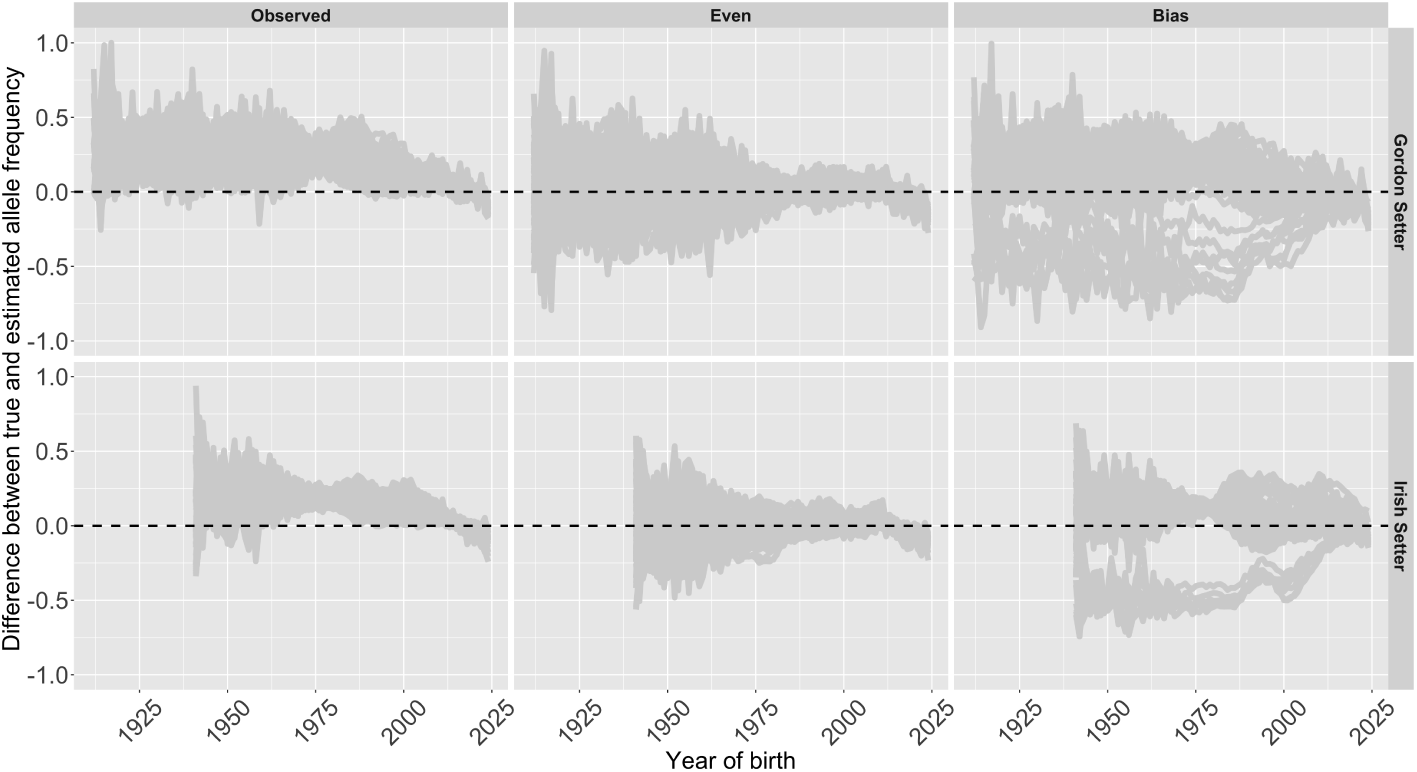
The difference between the true and estimated allele frequency (true – estimate) over the year of birth for the three simulated scenarios of distributing observed genotypes (“Observed”, “Even”, and “ Bias”) across the Gordon Setter and Irish Setter. The cloud of grey lines represents each of the 100 simulation replicates.

**Figure 6.**
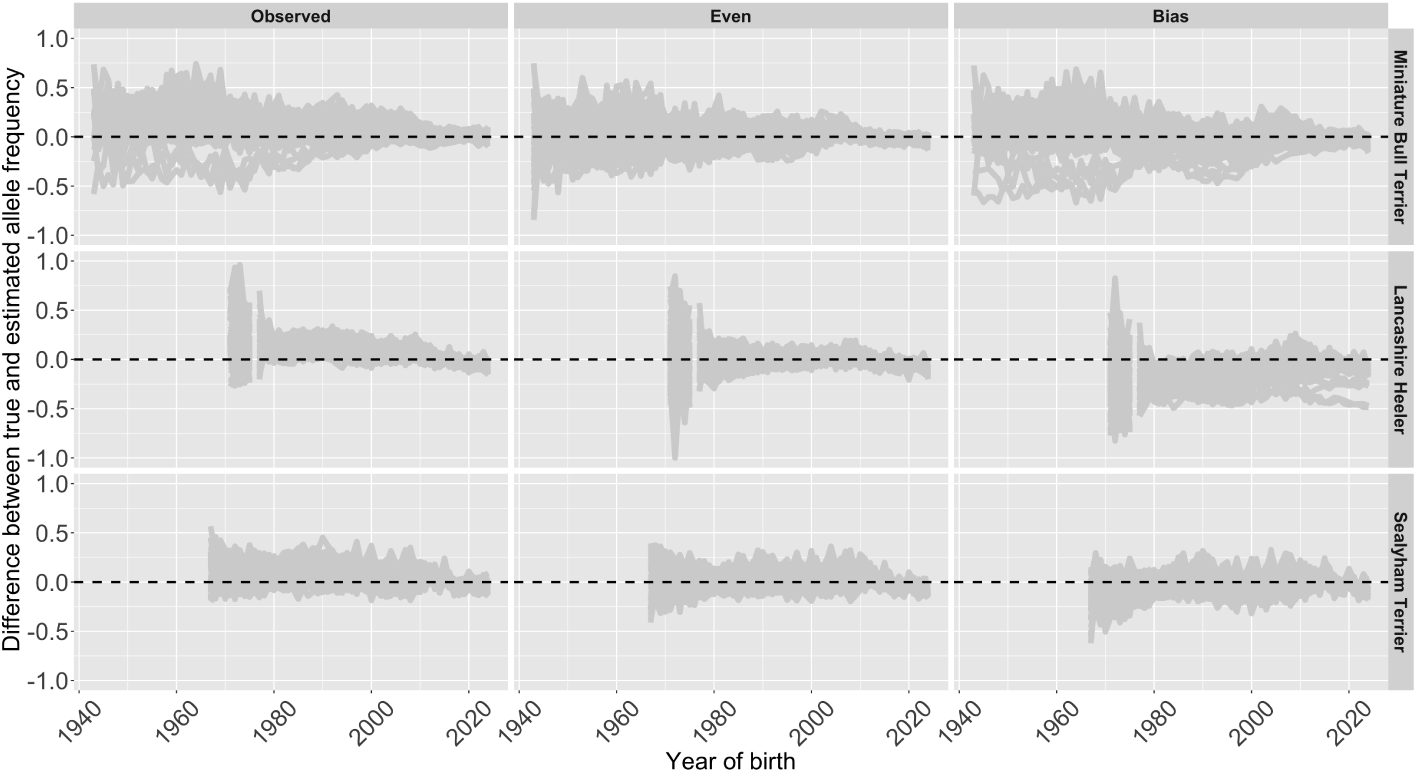
The difference between the true and estimated allele frequency (true – estimate) over the year of birth for the three simulated scenarios of distributing observed genotypes (“Observed”, “Even”, and “Bias”) across the Miniature Bull Terrier, Lancashire Heeler, and Sealyham Terrier. The cloud of grey lines represents each of the 100 simulation replicates.

For the “Observed” scenario in the Gordon Setter and the Irish Setter (Figure 5), between 1970 and 1985, there was a consistent disagreement between the estimated and true allele frequency in all 100 replicates. This was not observed in the “Even” scenario for either breed, where, for all years, the differences between “Observed-Even” were significant (p-values *<* 1.0 *×* 10*^−^*^11^). However, albeit minimally, we observed similar disagreement in the “Bias” scenario of the Irish Setter. This suggests that there was a high prevalence of lineages between 1970 and 1985 that did not undergo genetic testing in later years, which in the Irish Setter was possibly partly due to sampling or ascertainment bias. But, further research would be needed to confirm this. For all pedigrees, we observed a decreasing effect size towards zero from 1985 onward (not shown), with low to no significant difference between “Observed” and “Even” scenarios in the Miniature Bull Terrier, Lancashire Heeler, and Sealyham Terrier (2.6 *×* 10*^−^*^66^ *<* p-values *<* 6.7 *×* 10*^−^*^1^). From 2018, in all pedigrees except the Miniature Bull Terrier, a higher proportion of simulated repeats had a negative difference between true and estimated allele frequency in both the “Observed” and “Even” scenarios (Figure 5 and 6). Thus, it suggests that recent estimates in disease-causing allele frequency may be overestimated, but not due to the distribution of genotypes, since we also observed this in the optimal “Even” scenario.

Introducing bias for the distribution of genotypes had the largest influence on the two Setters (Figure 5). In the Irish Setter between 1960 and 1990, the simulations were separated into two groups: six simulations overestimated the allele frequency, the remaining 94 tended to underestimate the allele frequency. After genetic test availability, both groups converged. This was likely due to high relatedness from popular sires where the high random genetic drift was only partially captured by the later distributed genotypes [6, 58, 59]. Hence, illustrating the interaction between pedigree structure and distribution of genotypes. A similar grouping was observed in the Gordon Setter, however it was less distinct due to lower random genetic drift, assumed from the higher effective population size compared to the Irish Setters [6]. In the other breeds (Figure 6), especially the Lancashire Heeler, there was a higher likelihood of overestimating the allele frequency in the “Bias” scenario compared to the “Even” and “Observed” scenarios. The “Bias” scenario likely included more genotyped affected individuals than observed in the real data, leading to higher estimated base population allele frequencies for each UPG. Pairwise comparisons of the “Observed-Bias” scenario, showed significant difference across all years in the Gordon Setter and Lancashire Heeler (p-values *<* 4.4 *×* 10*^−^*^4^), suggesting that the real distribution of genetic tests differed from the “Bias” scenario in these breeds. Across the Miniature Bull Terrier, Irish Setter, and Sealyham Terrier there are periods of no significant difference. In the Irish Setter, this extended from 2015 to 2023, indicating potential presence of sampling and ascertainment biases in the real data influencing the performance of the probabilistic pedigree-based inference. Yet, with the suggested similarity of the “Bias” and “Even” scenario with many insignificant p-values in the “Bias-Even” pairwise comparison between 2012 and 2024 in the Irish Setter, these biases may not always reduce the quality of present day estimates.

Overall, these simulations highlight differences in the estimated allele frequencies from the distribution of genetic tests across different pedigree structures and distribution of genetic tests across pedigrees. The actual distribution of the genetic tests aligns more closely with the even distribution than the bias scenario. In recent years, all three scenarios were likely to overestimate the disease-causing allele frequency compared to the true population frequency per the simulation, suggesting that recent estimates were also limited by the number of genetic test results.

## Discussion

This study has evaluated probabilistic inference with unknown parent groups (UPG) at a single locus to propagate genotype information sourced from available genetic tests for two monogenic recessive eye conditions (PLL and PRA-rcd4) to all individuals in five Royal Kennel Club, UK (RKC) pedigrees with up to 74,000 records. The results showed estimates of disease-causing allele frequency in the years before and after the introduction of genetic testing, compared to when using only available genetic test results and hereditary status. Measured with leave-one-out cross-validation, the probabilistic pedigree-based inference achieved moderate to high accuracies across the five breeds. Further model-based validation using simulation highlights the influence of potential bias and the distribution of genotype data on the quality of the estimates, particularly for historical trends before the 2000s. In the following, we discuss the impact of the UPG assignment, the quality of the disease-causing allele frequency estimates and their interpretation, and suggest appropriate application of probabilistic pedigree-based inference for the RKC and other pedigree registration bodies.

### Little difference between UPG assignments

We considered nine different UPG assignments to address the inevitable missing pedigree data. Comparisons to models without UPGs identified subtle differences in allele frequency trends over the whole pedigree. Any observed differences typically occurred in the early pedigree, and this was expected since these individuals were far from genetic testing. Consequently, they have less information and were more influenced by how pedigree information was managed. Gengler *et al.* [35] reported a larger difference in estimated trends for the Myostatin gene with UPGs than without. They assigned UPG by year of birth across two sub-populations (beef and dual purpose). Thus, there was likely low genetic relatedness and high variance between the assigned UPG’s descendants due to the different selection goals resulting in different allele frequencies. In contrast, in dog pedigrees, we expect high genetic relatedness and low variance between UPG’s descendants; consequently, we only observed a small difference when including UPGs to without.

Overall, we chose the UPGYC100 assignment as the most appropriate pedigree handling in the current context of RKC-registered dogs. UPGYC100 grouped founders by year of birth and country of origin, while requiring a minimum of 100 “pseudorecords” in each UPG. This represented the changes in the allele frequency over time in the founders from random genetic drift and selection, and the differences between countries of origin due to geographical isolation[15, 60]. The number of imported dogs within RKC-registered pedigrees has been increasing since the easing of international dog transportation regulations in EU countries [2]. However, importation numbers had declined following the global COVID-19 pandemic, with more recent declines potentially associated with Brexit, although this relationship has not been established. Efforts are being made to re-establish these regulations to encourage importation to address low genetic diversity in some breeds [2]. A limitation in this data is that recorded country of origin may reflect country of export into the UK, rather than country of birth. Nevertheless, both year and country of origin are important grouping factors. Other grouping strategies were considered, including dog’s purpose (pet, show, and/or working); however, they could not be implemented due to data constraints.

Probabilistic inference shows a slower decline in the disease-causing allele frequency after genetic test introduction than estimates based on genetic tests and hereditary status

With probabilistic pedigree-based inference, the population trends for the disease-causing allele can be tracked across all individuals in the pedigree, instead of only the proportion of the pedigree with either a genetic test result or hereditary status. This not only offers more comprehensive monitoring of the disease-causing allele in recent years (see Figure 3), but also enables retrospective estimates within the early pedigree.

Prior to genetic test introduction, our estimates using genotype probabilities generally reported a lower disease-causing allele frequency than those using genetic test results and hereditary status. This was expected due to the known sampling bias present in this period for affected dogs to identify the disease-causing allele. In the 2000s, the Gordon Setter had a higher PRA-rcd4-causing allele frequency than the Irish Setter, in agreement with Downs *et al.* [40]. For the PLL-causing allele, our results reported that the Sealyham Terrier had the highest frequency, followed by the Lancashire Heeler, and then the Miniature Bull Terrier, prior to genetic test introduction. These estimates were broadly consistent with those reported by Gould *et al.* [11] for UK based Sealyham Terriers and Lancashire Heelers, although the allele frequencies estimated by [11] tended to be higher than those estimated here. In contrast, Gould *et al.* [11] estimated a PLL-causing allele frequency of 0.27 to 0.29 in Miniature Bull Terriers living in the UK, nearly double our estimates, whether using genetic tests and hereditary status only or the estimated genotype probabilities. For Gould *et al.* [11]’s 0.27 cohort, all dogs had unique sires and dams, and were not part of the original research population used by Farias *et al.* [39]. Yet Gould *et al.* [11] estimated allele frequency was possibly an overestimation due to participation and referral bias where owners with known or suspected risks were more likely to submit samples at the time. Conversely, the “Observed” scenario of Figure 6 suggests a tendency to underestimate the disease-causing allele frequency compared to the true defined by the simulation in the 2000s. Thus, the true estimate of disease-causing allele frequency before genetic test introduction likely falls between our estimate with probabilistic pedigree-based inference of 0.15 and Gould *et al.* [11] estimate of 0.27 for PLL in RKC-registered Miniature Bull Terriers.

After the genetic test introduction, for all five breeds, our estimates using genotype probabilities provide a higher disease-causing allele frequency compared to those based on genetic test results and hereditary status. This was expected due to two recognised biases. Firstly, there is the ascertainment bias, where breeders preferentially mate clear dogs to produce clear puppies. Secondly, there is inherent bias from the assignment of hereditary status, which is restricted to offspring of homozygous parents only [17]. In practice, this is limited to clear by clear matings, as matings involving homozygous affected dogs are extremely rare. In contrast, when using genotype probabilities, litters from clear by carrier matings are captured. Such matings are consistent with breed-management strategies that aim to reduce the production of offspring homozygous for the disease-causing allele, while avoiding removal of asymptomatic carriers and associated loss of genetic diversity, and as such can be expected to be commonly used. However, each puppy born in such litter has a 50% chance of carrying one copy of the disease-causing allele, contributing to the higher estimated disease-causing allele frequency compared with estimates based on genetic test result and hereditary status only. By considering the whole pedigree, both biases were likely reduced. Thus, as hypothesized, with probabilistic pedigree-based inference we observe a smaller decline in the disease-causing allele frequency around genetic test introduction.

### Quality of inference varies across the five pedigrees

The overall inference accuracy across the five breeds ranged from 0.47 (Miniature Bull Terriers) to 0.67 (Gordon Setters), which was broadly comparable to other studies examining single-locus inheritance, including Gengler *et al.* [35]. Although, direct comparisons are limited by differences in species, inference methods, validation designs, and performance metrics. Smaller pedigrees often require proportionally more genetic testing to achieve the same inference accuracies [e.g., 28]. Despite being the smallest two pedigrees, the Lancashire Heeler and the Sealyham Terrier have moderate accuracies, greater than the Miniature Bull Terrier, owing to the high number of genetically tested dogs (see Figure 1). In contrast, the Miniature Bull Terrier’s inference quality is limited by the relatively steep annual decline in the number of genetically tested dogs following the genetic test publication. Further limitations include the relatively high pedigree incompleteness around genetic test introduction (see Supplementary Figure S1) and high effective population size [6], together with the lowest disease-causing allele frequency (see Table 5) [33, 34]. For rare disease-causing alleles (frequency less than 5%), alternative techniques to probabilistic pedigree-based inference exist, such as ancestral recombination graphs [61]. However, this requires multi-locus genotype data rather than a few single-locus genotypes, as pedigree dogs and other small pedigrees are currently routinely using, though multi-locus genotype is now becoming increasingly used too [41]. Nonetheless, Table 5 shows that moderate to high accuracies were achieved using probabilistic pedigree-based inference with disease-causing allele frequencies ranging from 0.20 to 0.27 across the two eye conditions and five breeds.

Another indication of inference quality can be observed from the “Observed” genotyping scenario using the simulated data in Figures 5 and 6. Unlike the leave-one-out cross-validation, this considered the whole pedigree time-frame, compared to just those with genetic test data. The two validation approaches provide complementary insights into potential bias in estimated disease-causing allele frequencies. The genotype-specific mean signed dosage errors from the leave-one-out cross-validation indicated small systematic overestimation of allele frequency in the Miniature Bull Terrier and the Irish Setter, with insufficient evidence for the other three breeds. Nonetheless, the maximum absolute differences attributed to these errors was only 0.018, suggesting very little systematic bias within the genetically tested cohort. For all pedigrees, except the Miniature Bull Terrier, the simulated data showed a higher likelihood of overestimating than underestimating the disease-causing allele from 2018 onward through its stochastic repeats. This can be explained by two reasons. First is the potential selection within the simulation that is not captured by the available genetic test results in the genotype file. In the “Even” genotyping scenario, the distribution of genotypes captures less of the selection, resulting in higher overestimates in allele frequencies. In the “Observed” genotyping scenario the effect is minimal, however, indicates potential error in the assignment of hereditary status, as the intensity of selection in the simulation is based on available genetic tests and hereditary status. These errors could include the assignment of hereditary status to offspring with unrecorded genotype in one or both parents or due to pedigree error reducing the certainty in the estimated genotype probabilities. This is not surprising given an expected pedigree error between 1 to 9% in dogs [21, 22]. Second, particularly in the Setters, the number of RKC registrations per year has been declining. Consequently, while the proportion of genetically tested dogs in the Setters remains consistent (see Figure 1), the number of genetic test results has been declining. Thus, it reduces the quality of the estimated disease-causing allele frequencies [e.g., 28]. This overestimation was not observed in the “Observed” genotyping scenario of the Miniature Bull Terrier, likely due to a relatively high decline in hereditary status assignment (see Figure 1). While this may imply the use of carrier by clear matings, it more likely indicates increased matings where one or both parents have unknown genotypes, particularly as the breed has become more popular and is crossed with standard Bull Terriers (where the PLL-causing allele is very rare, thus untested) through the established crossbreeding scheme [11, 62, 63]. Regardless, our finding supports Lewis [22]’s recommendation to restrict hereditary status to two generations after genetic testing, but yet to be implemented by the RKC. Moreover, this highlights the need for continuous genetic testing of dogs in addition to performing probabilistic pedigree-based inference for population monitoring of disease-causing alleles.

### Application of probabilistic inference in pedigreed populations

Primarily, this study aims to improve population monitoring of disease-causing alleles in small pedigreed populations. Additionally, the genotype probabilities for individuals can inform untested dogs’ genotypes, but high accuracy is essential. Van den Berg *et al.* [31] reported high accuracy for assignment of carrier status in dairy cattle using the F1 score, but in dogs we found the accuracy was moderate for carriers (around 0.5) and low for affected (around 0.3), as presented through the calibrated F1-scores in Table 6. Although, differences in the performance metrics used limit direct comparisons between van den Berg *et al.* [31]’s and the present study. Exceptions exist, such as historic sires with data from many relatives, where genotypes can be assigned with high certainty [64, 65]. Alternatively, the RKC could use genotype probabilities to identify “at-risk” dogs for genetic testing as described in the second additional file of the supplementary materials. However, labelling dogs as “at-risk” could discourage testing due to negative perceptions, even if dog owners consented [19, 66, 67]. Without consent, the disclosure of suspected health risks to unsuspecting dog owners raises ethical concerns, particularly given established guidance on communication of secondary genetic findings established in humans [68]. Furthermore, with the moderate to low accuracies and need for stringent thresholds, there may be many false positives, consequently identifying “at-risk” dogs may not provide sufficient benefit compared to promoting genetic testing for all dogs. Alternatively, genotype probabilities across multiple loci could be combined into a score and mating plans can be devised to minimize the probability of affected genotypes, while tracking carrier genotypes. While it is tempting to strictly remove mating of carrier genotypes, this only creates additional bottlenecks in dog pedigrees which are typically already limited through closed registries, popular sires, historical bottlenecks and selection for a range of traits [6, 7, 20], so more active management is warranted and estimated genotype probabilities provide information for such a system.

Overall, the application of probabilistic pedigree-based inference is appropriate mostly for population-level analysis and will provide more comprehensive monitoring of disease-causing alleles in pedigreed populations compared to relying solely on the proportion of the pedigree with genotype information. In addition to disease-causing allele frequencies, the genotype frequencies can be monitored overtime, whether phased or unphased. Thus, informing whether the recessive disease-causing allele is maintained predominantly among heterozygous (carrier) dogs or increasing the expected frequency of homozygous genotypes (affected). Such information could help inform breeding strategies that seek to reduce the occurrence of recessive hereditary diseases whilst conserving genetic diversity; an important priority in many UK dog breeds [19, 69]. However, given variation in genotype inference quality among breeds and genotype classes, estimated genotype frequency trends should be interpreted as indicative rather than definitive, alongside the uncertainty in the underlying estimates. Nevertheless, the use of estimated genotype probabilities for population trends will better inform the RKC for breed-specific recommendations to reduce the future prevalence of these diseases; however, there are two anticipated challenges in the application. First, there is the potential for increased insurance premiums or exclusions for specific breeds and hereditary conditions [19, 70]. Such changes could make adequate insurance less accessible or affordable for some dog owners, potentially increasing the number of uninsured or underinsured dogs and risk of delayed veterinary treatment [71]. The cost-of-living crisis has already placed additional financial pressure on dog owners, likely contributing to reduced uptake or retention of pet insurance and an increase in the number of dogs being surrendered because owners can no longer afford their care [72–74]. Second, dog owners may not opt for genetic testing due to demotivation from the perceived ineffectiveness, as the frequency is higher than previously thought, or because they do not see the need with the estimated genotype probabilities. Yet, the reason the disease-causing allele frequency is higher is likely due to the proportion of the pedigree that does not use genetic testing and the capturing of clear by carrier matings which produce litters without homozygous affected offspring whilst minimising loss in genetic diversity. Moreover, the quality of the probabilistic pedigree-based inference relies upon the continued genetic testing of dogs. Therefore, to capture the advantages of probabilistic inference, it is imperative to sustain educational initiatives and promotional campaigns for genetic testing to dog owners.

### Study limitations

There are three key limitations to this study. Firstly, when assigning dummy individuals to individuals with only one known parent, we ignored the potential of multiple sire litters when considering littermates. Multiple sire litters are possible in dogs, although the frequency of illegitimate multi-sires is unknown [75]. Additionally it is possible that dogs sharing a known sire and date of birth may have different dams and belong to different litters. However, with such a small number of half-founders (see Supplementary Table S1), changing the strategy used because of possible false full sibling assignment is unlikely to make any difference to the estimates.

Secondly, the model assumed no relationship between UPGs, which is unlikely. Metafounders are UPGs that consider the potential relationship between and within each grouping. Although introduced with multi-loci genotype data [76], other studies have highlighted the potential to infer UPG relationships using the rate of inbreeding over time [77]. However, it assumes a consistent rate of inbreeding over time, which is not true for many dog breeds [6].

Finally, despite the leave-one-out cross-validation being implemented to reduce the influence of missing data, the approach still presents limitations for two reasons. One, we assume the genetic test results to be error-free. However, errors are possible, particularly during sampling, damage during postage, or laboratory error [78, 79]. Falsification is also possible, although thought to be rare [17, 69]. In the same way, largely pedigree is assumed error free which is unlikely the case. The second concerns the potential underestimation of the affected calibrated F1 score. The majority of genetically tested “affected” dogs were tested in the 2000s as part of the initial identification of the disease-causing allele, with no ancestors genetically tested and few descendants or other relatives. Consequently, the estimated genotype probabilities will rely on the prior that reports a low disease-causing allele frequency. Hence, there is a lower likelihood of correctly identifying these individuals. For present day estimates of “affected” individuals, the calibrated F1-score may be higher with the accumulation of genetic tests over generations. Although this could lead to an underestimation in the overall inference accuracy, this is likely counteracted by the possible overestimation from the within family clustering of genetic tests after genetic test introduction. Moreover, since the leave-one-out cross-validation was restricted to genetically tested individuals, a potentially better-connected subset of the pedigree, the reported performance may not generalise to historic, untested, or poorly connected dogs, where the overall inference accuracy would likely be lower.

## Conclusion

By combining the available genetic tests and pedigree information, this study demonstrates a method to more comprehensively monitor the monogenic disease-causing allele frequency in pedigreed populations than relying on available genetic tests and hereditary status alone. The quality of the inference depends on the frequency of the disease-causing allele, the number and distribution of genetic tests, and the yearly census size of the pedigree. Thus, we show that the inference quality is inconsistent across pedigrees for monogenic recessive hereditary diseases. The assignment of genotype to non-genotyped dogs or identification of “at-risk” dogs for genetic testing based on the estimated genotype probabilities was considered, but not recommended due to the moderate to low accuracies for carrier or affected status in these pedigrees. However, such a strategy may be effective in the case of popular sires, for example. Future studies will explore the use of these estimated genotype probabilities in downstream analyses to understand the key genetic processes that drive these changes in disease-causing allele frequencies. Moreover, as the uptake of multi-locus genotype data increases in dogs, the quality of probabilistic pedigree-based inference is expected to improve.

## Ethics approval and consent to participate

Not applicable

## Consent for publication

Not applicable

## Availability of data and materials

The data for this study were provided by the Royal Kennel Club in the UK. The data can be requested for research purposes through the Royal Kennel Club. All scripts used, across R and Python, were provided through the GitHub repository: https://github.com/HighlanderLab/RCraddock_AlleleFetch_Geno.

## Competing interests

JJI is employed by The Royal Kennel Club, which provided the pedigree and genetic testing data used in this study. CM is an employee of an institution that offers canine genetic testing as a commercial service. The authors declare no other competing interests.

## Funding

The authors acknowledge funding from the Kennel Club Charitable Trust and the BBSRC ISP to The Roslin Institute (BBS/E/RL/230001A and BBS/E/RL/230001C), and support from the University of Edinburgh.

For the purpose of open access, the author has applied a CC BY public copyright licence to any Author Accepted Manuscript version arising from this submission.

## Author’s contributions

RFC performed all the analyses and drafted the manuscript, MJF contributed to the initial project design, CM and JJI contributed to the interpretation of results, PW and SS co-supervised the study, GG designed and supervised the study. All authors read, edited, and approved the final version of the manuscript.

## Acknowledgements

We are grateful to Audrey Martin, Xinger Tang, Jeffrey Schoenebeck, and Pau Navarro as well as members of the Highlander Lab for their valuable input into the project.

## Additional Files

### Additional file 1 — Supplementary tables and figures

#### Comparisons of pedigree structure across breeds

Table S1 provides the relevant background on the pedigree structure for all five breeds studied, considering the potential influence on inference quality. The Lancashire Heeler has the lowest pedigree depth and is the youngest pedigree by birth year. The Miniature Bull Terrier has the highest pedigree depth despite being only the third oldest pedigree by year of birth, behind Gordon Setter and Irish Setter. With many dogs imported, the Sealyham Terrier has the highest percentage of founders, over 2.5 times more than the second highest pedigree. The total percentage of half-founders is small, ranging from 0.19 to 0.71%. For all five pedigrees, there is a lower proportion of unique sires than dams by about half.

#### Pedigree completeness across birth cohorts

Pedigree completeness was assessed to examine the distribution of missing ancestral information across the recorded pedigree timelines and to illustrate that dogs with unknown parentage (pedigree founders) are not necessarily restricted to the earliest birth cohorts.

For each dog, the proportion of known ancestral positions was calculated separately for each of the first five ancestral generations where generation 1 comprised parents, generation 2 grandparents, with subsequent generations representing progressively more distant ancestors. At generation g, completeness was calculated as the number of known ancestral positions divided by the maximum possible number of positions (2*^g^*). An ancestor was considered known if their individual identity was recorded. For example, if only one parent is known, generation 1 would have 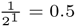, so 50% completeness.

Completeness values were then averaged across dogs sharing the same birth year (using the inferred dates of birth), separately by ancestral generation and breed as presented in Supplementary Figure S1.

#### Quantification of uncertainty in estimated allele and genotype frequencies

To quantify the model-based uncertainty in the estimated allele and genotype frequencies, a simple Monte Carlo approach was used with 10,000 repeats. In each repeat, first a discrete genotype was sampled for each individual based on the estimated genotype probabilities from AlphaPeel’s output using UPGYC100 pedigree information and all available genetic test results; second, the allele and genotype frequencies were estimated for each birth year.

Once all results from each repeat were collected, the 2.5% and 97.5% quantiles in each year were used for the lower and upper range of the approximate 95% credible intervals and plotted through the ribbons in Supplementary Figures S2 and S3.

Supplementary Figures S2 and S3 show that uncertainty was highest prior to 1980. For example, the Lancashire Heeler 95% credible interval covered the whole range of 0 to 1 in 1970 to 1972. As sample sizes increased with more pedigree information from digitisation and individuals were born closer to genetic test introduction in the late 2000s, the credible intervals reduced in size for all five pedigrees. Some uncertainty remains in the allele frequencies of the Sealyham Terrier and Lancashire Heeler from 2000s onwards whereas there is no observable uncertainty in the other three pedigrees (Supplementary Figure S2). The Sealyham Terrier and the Lancashire Heeler were the two pedigrees with the smallest census size, thus the sustained small uncertainty through genetic test introduction is likely a factor of the limited sample size.

As not all individuals recorded in the pedigree have a recorded or inferred date of birth, Supplementary Figure S4 shows the disease-causing allele frequency trends with uncertainty by generation, where a generation of 1 represents all the pedigree founders. As all dogs in the pedigree were allocated a generation number, all individuals recorded in the pedigree are represented in this plot.

The simple Monte Carlo approach used to quantify the uncertainty in the estimated allele and genotype frequencies independently sampled genotypes. Thus, covariance among genotypes, such as that arising from pedigree, were not accounted for. By ignoring the covariance amongst genotypes and hence the uncertainty propagated through the pedigree, the uncertainty in allele and genotype frequencies may be underestimated since related individuals contribute less independent information than unrelated. Alternatively, ignoring the covariance may overestimate the uncertainty due to pedigree providing inheritance constraints. Future work would account for the covariance between genotypes by sampling from the estimated genotype and transmission probabilities conditional on defined family beliefs. Although the change in the credible intervals would likely only be marginal, due to the relatedness, wider intervals would be expected.

#### Mean error of allele frequencies from leave-one-out cross-validation

In Table 6, the genotype-specific mean signed error dosage were presented for each breed through the leave-one-out cross-validation. Although this highlighted the shrinkage towards intermediates with overestimation of clear genotypes and underestimation of carrier and affected genotypes of the genetically tested cohort, how this influences the overall estimated disease-causing frequencies depends on the distribution of genotype samples. Supplementary Table S2 presents the mean signed error for the allele frequencies of each breed. The sample size, n, was multiplied by the mean signed dosage error then summed across the three genotype states. These were then divided by two times the total sample size across the three genotype states.

**Table S1.**
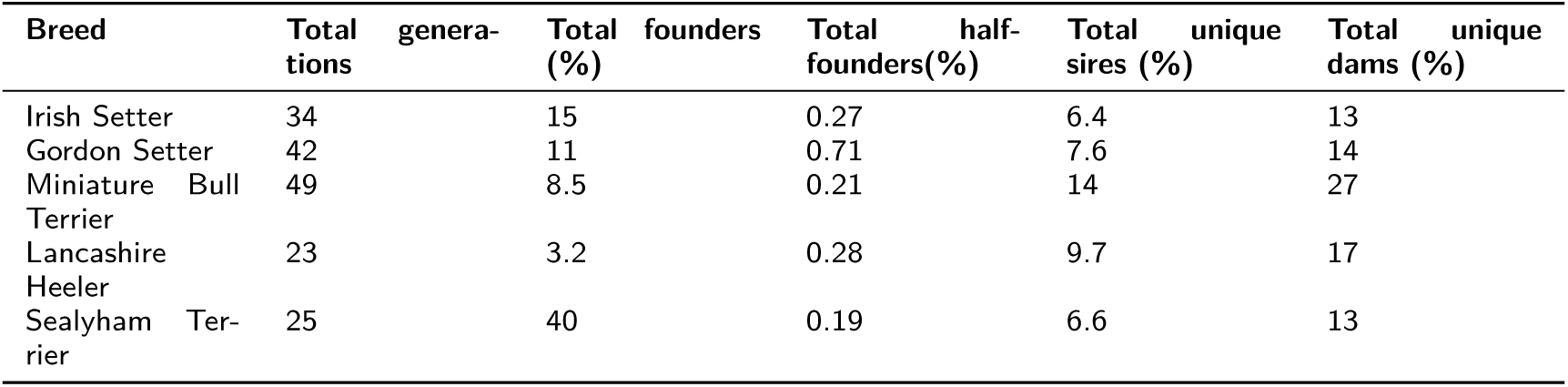
Summary of pedigree structure of the five breeds.

| Breed | Total generations | Total founders (%) | Total half-founders (%) | Total unique sires (%) | Total unique dams (%) |
| --- | --- | --- | --- | --- | --- |
| Irish Setter | 34 | 15 | 0.27 | 6.4 | 13 |
| Gordon Setter | 42 | 11 | 0.71 | 7.6 | 14 |
| Miniature Bull Terrier | 49 | 8.5 | 0.21 | 14 | 27 |
| Lancashire Heeler | 23 | 3.2 | 0.28 | 9.7 | 17 |
| Sealyham Terrier | 25 | 40 | 0.19 | 6.6 | 13 |

**Table S2.** Mean signed error for the allele frequencies of each breed based on the genotype-specific mean errors and number of genetically tested dogs in each genotype class.

| Breed | Mean signed error disease-causing allele frequency (95%CI) |
| --- | --- |
| Gordon Setter | -0.008 (-0.023, +0.007) |
| Irish setter | +0.010 (+0.000, +0.021) |
| Sealyham | +0.007 (-0.013, +0.027) |
| Lancashire Heeler | -0.0018 (-0.019, +0.015) |
| Miniature Bull Terrier | +0.018 (+0.002, +0.034) |

### Additional file 2 — Description of tool for identifying “at-risk” dogs for genetic testing

#### Suggested tool for identifying “at-risk” dogs for genetic testing

After performing the probabilistic pedigree-based inference, every dog in the pedigree has estimated genotype probabilities, one for each state (“clear”, “carrier”, and “affected”), where the sum of all the genotype probabilities equals 1 for each individual. For dogs that are not genetically tested, this could be used to identify “at-risk” dogs for genetic testing. For this, a Receiver Operating Characteristic (ROC) curve can be used. A ROC curve quantifies the trade-off between sensitivity and specificity for identifying at-risk dogs (either carrier or affected). The continuous scale was derived from the sum of the carrier and affected estimated genotype probabilities (noted as “non-clear”), ranging from 0 to 1. Varying thresholds were used to convert this continuous scale into a binary classification of “non-clear” (positive) or “clear” (negative), which was then compared to the true genotype collected during the leave-one-out cross-validation. By combining carrier and affected genotype probabilities, the true samples for “non-clear” and “clear” are more balanced (see Table 6). Additionally, this approach enables the identification of at-risk dogs for segregating the disease-causing allele. Supplementary Figure S5 presents the ROC curve showing the trade-off between correctly detecting non-clear dogs (true positive rate) and incorrectly classifying clear dogs as non-clear (false positive rate) at thresholds from 0 to 1. All pedigrees perform better than random assignment though far from perfect, with the Gordon Setter showing the highest performance, indicated by the curve being closest to the top left corner, consistent with overall inference accuracy, reliability, and calibrated F1-score.

When setting a threshold to identify at-risk dogs for allocation of finite resources (such as promotional material for genetic testing), there are many factors to consider by the breed registration. In addition to the quality of the inference, these include (but not limited to) the prevalence of the disease, the severity of the disease and potential treatment, and the rate of inbreeding in the breed. For example, a probability threshold of 0.45 or more may be used. Using the ROC curve for PRA-rcd4 in the Gordon Setter, this is estimated to include 60% of the pedigree sample, correctly identifying around 85% of all non-clear dogs. However, this means 15% are missed, and 40% of all clear dogs are incorrectly identified as non-clear. Overall, this would not be used to assign a “non-clear” status, but rather behind the scenes for the breed registration to direct resources to promote genetic testing to at-risk dogs.

**Figure S1.**
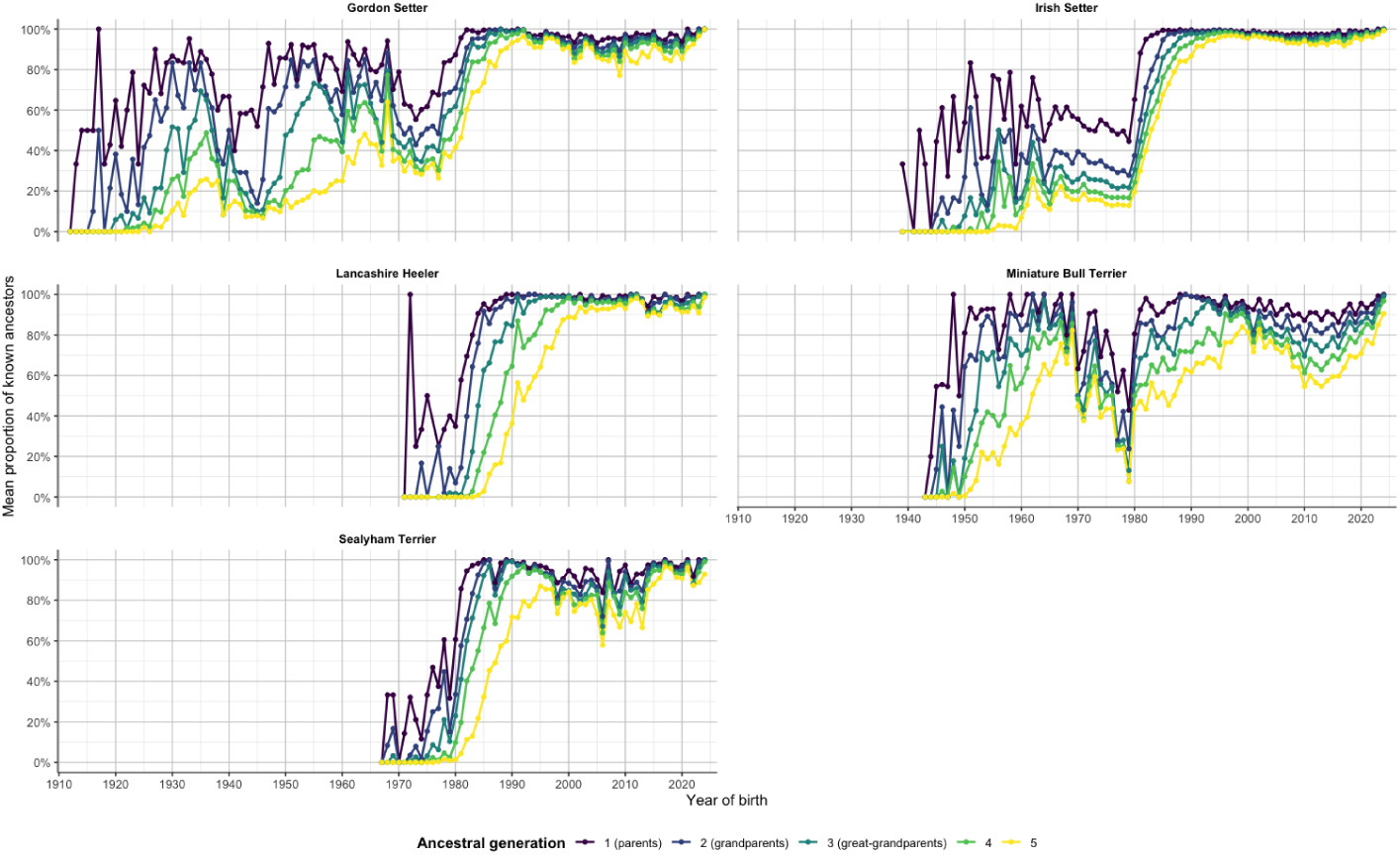
Mean proportion of known ancestral proportions at each of the first five generations by year of birth and breed. Lower completeness indicates a greater proportion of ancestral positions could not be assigned to individually identified dogs.

**Figure S2.**
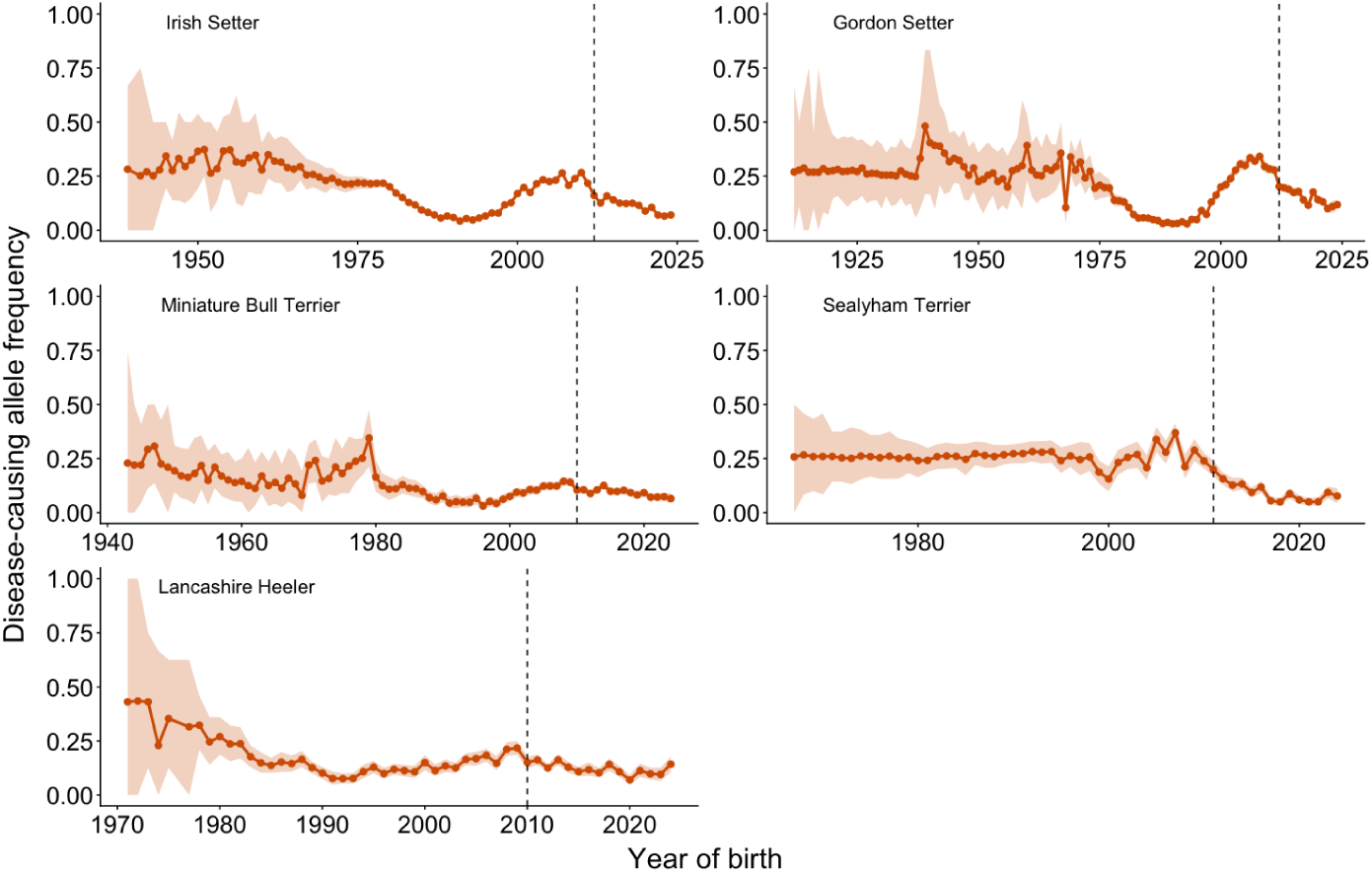
Disease-causing allele frequencies by year of birth with 95% credible intervals, representing the model-based uncertainty for PRA-rcd4 in Irish Setter and Gordon Setter, and PLL in the Miniature Bull Terrier, Sealyham Terrier, and Lancashire Heeler. Note the different scales on the x-axis due to differing timeframes across the pedigrees.

**Figure S3.**
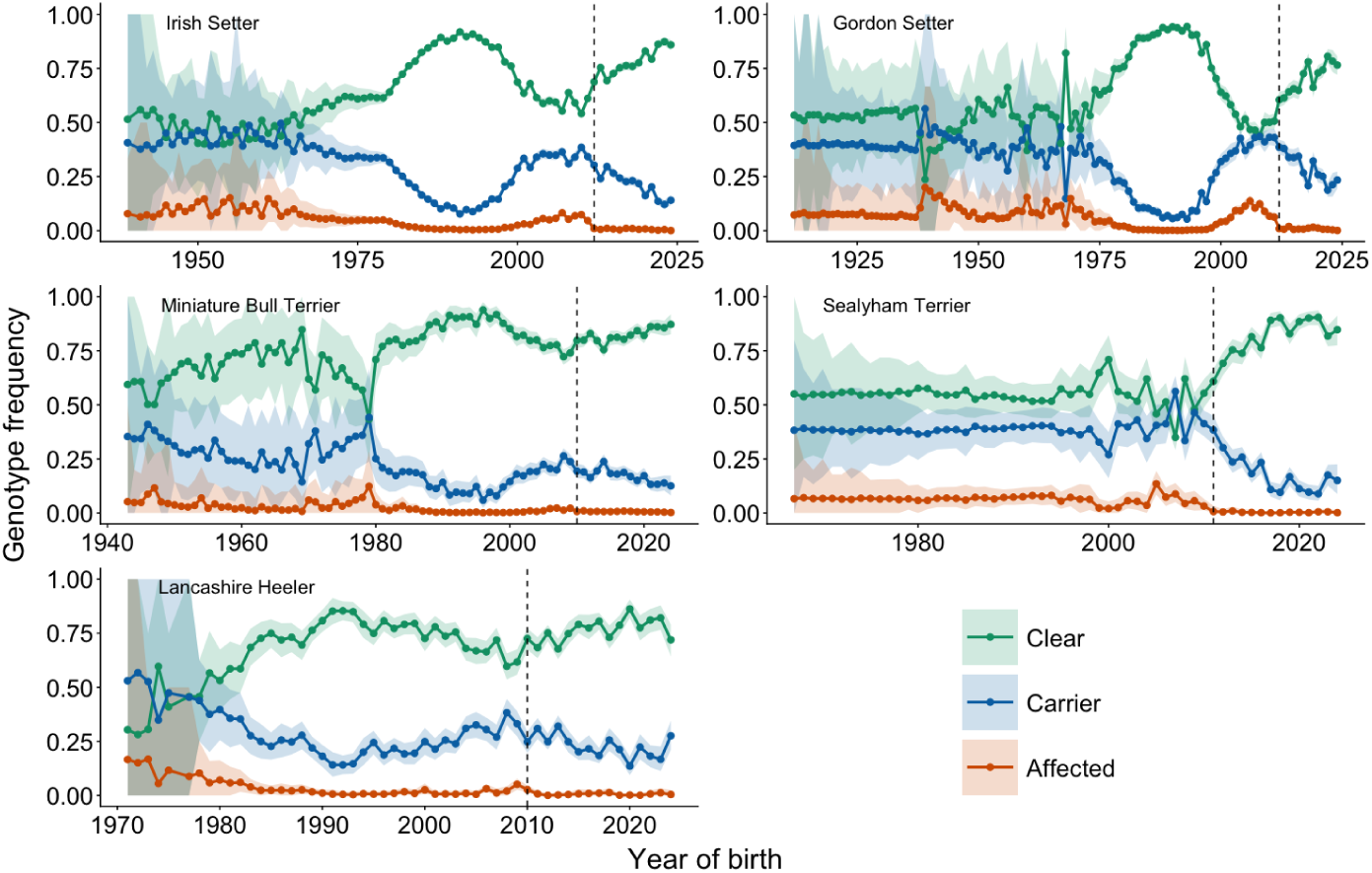
Genotype frequencies by year of birth with 95% credible intervals, representing the model-based uncertainty for PRA-rcd4 in Irish Setter and Gordon Setter, and PLL in the Miniature Bull Terrier, Sealyham Terrier, and Lancashire Heeler. Note the different scales on the x-axis due to differing timeframes across the pedigrees.

**Figure S4.**
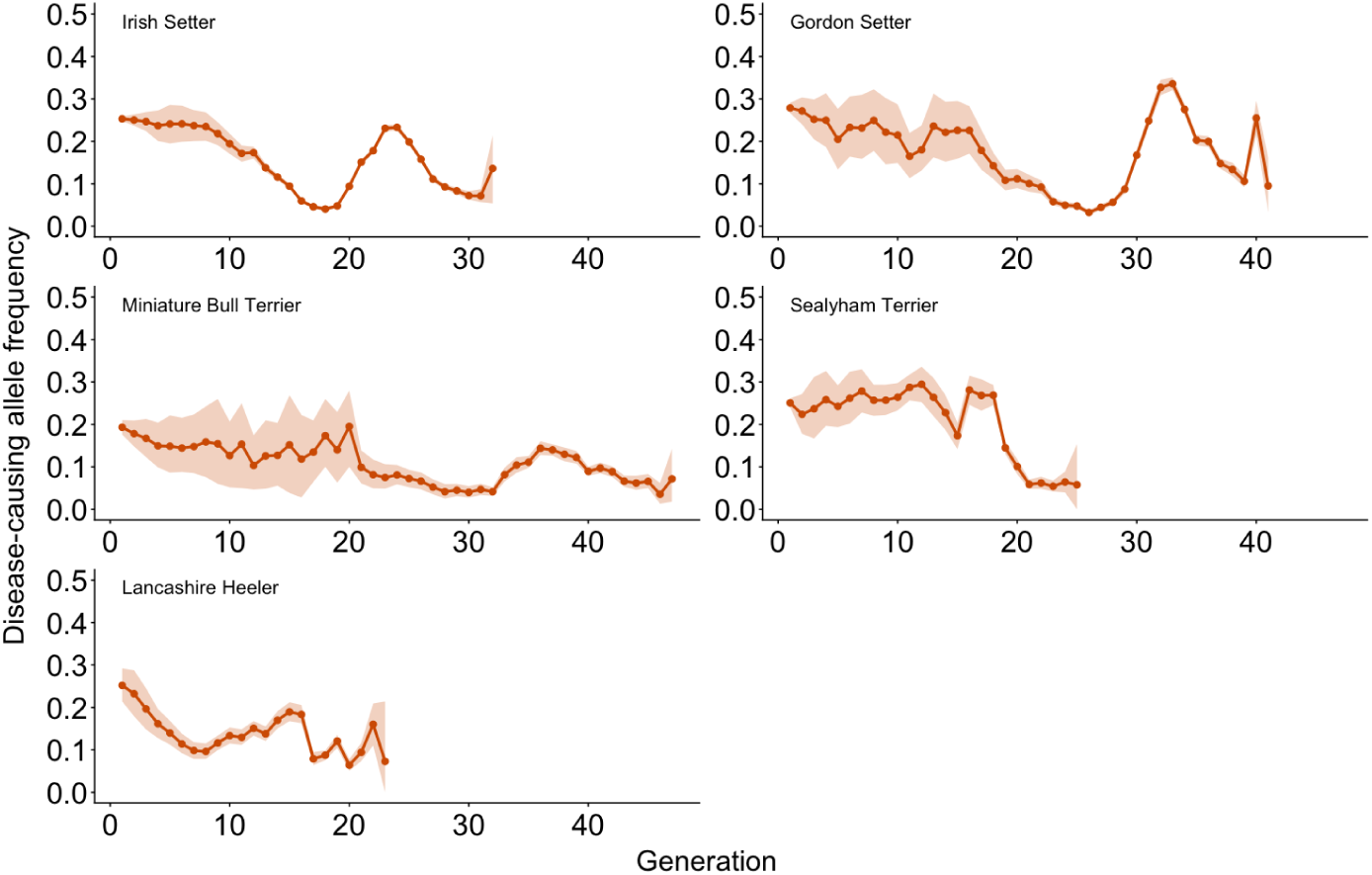
Disease-causing allele frequencies by generations with 95% credible intervals, representing the model-based uncertainty for PRA-rcd4 in Irish Setter and Gordon Setter, and PLL in the Miniature Bull Terrier, Sealyham Terrier, and Lancashire Heeler. Generation 1 contains all pedigree founders

**Figure S5.**
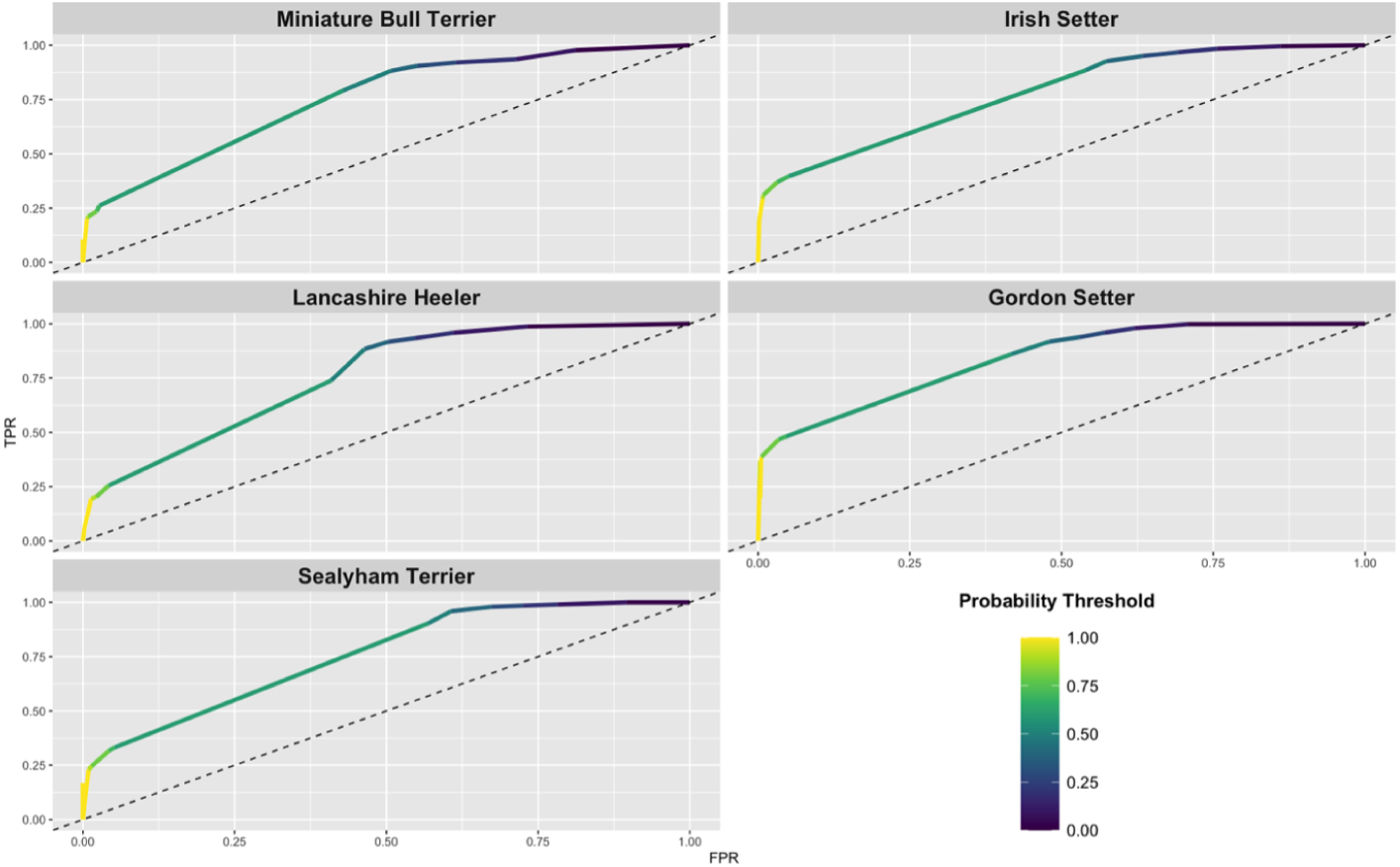
ROC curve for non-clear (carrier or affected) genotype assignment from the estimated genotype probability thresholds in all five breeds.

## References

1. Meuwissen T. Genetic management of small populations: A review. Acta Agriculturae Scandinavica, Section A — Animal Science. 2009;59(2):71–79.

2. Ilska JJ, Ryan P, Tolhurst D. Modern pedigree dogs in the United Kingdom Kennel Club: a journey through shifting population landscapes and demography. Companion Animal Health and Genetics. 2025;12(1):3.

3. Donner J, Anderson H, Davison S, Hughes AM, Bouirmane J, Lindqvist J, et al. Frequency and distribution of 152 genetic disease variants in over 100,000 mixed breed and purebred dogs. PLoS genetics. 2018;14(4):e1007361.

4. Donner J, Freyer J, Davison S, Anderson H, Blades M, Honkanen L, et al. Genetic prevalence and clinical relevance of canine Mendelian disease variants in over one million dogs. PLoS Genetics. 2023;19(2):e1010651.

5. Leroy G. Genetic diversity, inbreeding and breeding practices in dogs: results from pedigree analyses. The Veterinary Journal. 2011;189(2):177–182.

6. Lewis T, Abhayaratne B, Blott S. Trends in genetic diversity for all Kennel Club registered pedigree dog breeds. Canine genetics and epidemiology. 2015;2(1):13.

7. Marsden CD, Ortega-Del Vecchyo D, O’Brien DP, Taylor JF, Ramirez O, Vilà C, et al. Bottlenecks and selective sweeps during domestication have increased deleterious genetic variation in dogs. Proceedings of the National Academy of Sciences. 2016;113(1):152–157.

8. Mooney JA, Yohannes A, Lohmueller KE. The impact of identity by descent on fitness and disease in dogs. Proceedings of the National Academy of Sciences. 2021;118(16):e2019116118.

9. Vasiliadis D, Metzger J, Distl O. Demographic assessment of the Dalmatian dog–effective population size, linkage disequilibrium and inbreeding coefficients. Canine medicine and genetics. 2020;7(1):3.

10. Tournebize R, Chu G, Moorjani P. Reconstructing the history of founder events using genome-wide patterns of allele sharing across individuals. PLoS Genetics. 2022;18(6):e1010243.

11. Gould D, Pettitt L, McLaughlin B, Holmes N, Forman O, Thomas A, et al. ADAMTS17 mutation associated with primary lens luxation is widespread among breeds. Veterinary Ophthalmology. 2011;14(6):378–384.

12. Gharahkhani P, O’Leary C, Duffy D, Bernays M, Kyaw-Tanner M, et al. Primary Lens Luxation in Australian Tenterfield and Miniature Bull Terriers is due to an old ADAMTS17 mutation and is an additive trait. Open Genomics J. 2012;5:7–13.

13. Gharahkhani P, O’Leary CA, Duffy DL, Kyaw-Tanner M. Potential modifying loci associated with primary lens luxation, pedal hyperkeratosis, and ocular phenotypes in Miniature Bull Terriers. Investigative Ophthalmology & Visual Science. 2015;56(13):8288–8296.

14. Nicholas FW, Tanmen I, Sydney Information Hub. OMIA:000588-9615: Online Mendelian Inheritance in Animals (OMIA) [dataset]. The University of Sydney; 2025. https://omia.org/OMIA000588/9615/. DOI: 10.25910/2AMR-PV70. Accessed 14 October 2026.

15. Parker HG, Dreger DL, Rimbault M, Davis BW, Mullen AB, Carpintero-Ramirez G, et al. Genomic analyses reveal the influence of geographic origin, migration, and hybridization on modern dog breed development. Cell reports. 2017;19(4):697–708.

16. British Veterinary Association, The Royal Kennel Club, International Sheep Dog Society. BVA/RKC/ISDS Eye Scheme – Inherited Eye Disease: Breeds affected and type of disease. British Veterinary Association; 2026. https://www.bva.co.uk/media/6771/known-inherited-ocular-diseases-kiod-2026.pdf. Accessed 14 October 2026.

17. Lewis TW, Mellersh CS. Changes in mutation frequency of eight Mendelian inherited disorders in eight pedigree dog populations following introduction of a commercial DNA test. PloS one. 2019;14(1):e0209864.

18. Rokhsar JL, Canino J, Raj K, Yuhnke S, Slutsky J, Giger U. Web resource on available DNA variant tests for hereditary diseases and genetic predispositions in dogs and cats: An Update. Human genetics. 2021;140(11):1505–1515.

19. Farrell LL, Schoenebeck JJ, Wiener P, Clements DN, Summers KM. The challenges of pedigree dog health: approaches to combating inherited disease. Canine genetics and epidemiology. 2015;2(1):3.

20. Leroy G, Baumung R. Mating practices and the dissemination of genetic disorders in domestic animals, based on the example of dog breeding. Animal Genetics. 2011;42(1):66–74.

21. Leroy G, Danchin-Burge C, Palhière I, Baumung R, Fritz S, Mériaux J, et al. An ABC estimate of pedigree error rate: application in dog, sheep and cattle breeds. Animal Genetics. 2012;43(3):309–314.

22. Lewis T. The impact of incorrectly recorded parentage on inferred genotypes over multiple generations. Canine Genetics and Epidemiology. 2019;6(1):6.

23. Elston RC, Stewart J. A general model for the genetic analysis of pedigree data. Human heredity. 1971;21(6):523–542.

24. Janss L, Van Arendonk J, Van der Werf J. Computing approximate monogenic model likelihoods in large pedigrees with loops. Genetics Selection Evolution. 1995;27(6):567–579.

25. Kerr R, Kinghorn B. An efficient algorithm for segregation analysis in large populations. Journal of Animal breeding and Genetics. 1996;113(1-6):457–469.

26. Whalen A, Ros-Freixedes R, Wilson DL, Gorjanc G, Hickey JM. Hybrid peeling for fast and accurate calling, phasing, and imputation with sequence data of any coverage in pedigrees. Genetics Selection Evolution. 2018;50(1):67.

27. Ros-Freixedes R, Whalen A, Chen CY, Gorjanc G, Herring WO, Mileham AJ, et al. Accuracy of whole-genome sequence imputation using hybrid peeling in large pedigreed livestock populations. Genetics Selection Evolution. 2020;52(1):17.

28. Ros-Freixedes R, Whalen A, Gorjanc G, Mileham AJ, Hickey JM. Evaluation of sequencing strategies for whole-genome imputation with hybrid peeling. Genetics Selection Evolution. 2020;52(1):18.

29. Fernández SA. An algorithm to sample genotypes in complex pedigrees. Iowa State University; 2001.

30. Fernández SA, Fernando RL, Guldbrandtsen B, Totir LR, Carriquiry AL. Sampling genotypes in large pedigrees with loops. Genetics Selection Evolution. 2001;33(4):337.

31. van den Berg I, Nguyen T, Nguyen T, Pryce J, Nieuwhof G, MacLeod I. Imputation accuracy and carrier frequency of deleterious recessive defects in Australian dairy cattle. Journal of Dairy Science. 2024;107(11):9591–9601.

32. Van Doormaal B, Kistemaker G. Managing genetic recessives in Canadian Holsteins. Interbull Bulletin. 2008;38:75–79.

33. Tier B, Henshall JM. Limits to genotypic probabilities for single loci. In: Application of New Genetic Technologies to Animal Breeding. BioOne; 2005. p. 366–370.

34. Ullah E, Mall R, Abbas MM, Kunji K, Nato AQ, Bensmail H, et al. Comparison and assessment of family-and population-based genotype imputation methods in large pedigrees. Genome research. 2019;29(1):125–134.

35. Gengler N, Mayeres P, Szydlowski M. A simple method to approximate gene content in large pedigree populations: application to the myostatin gene in dual-purpose Belgian Blue cattle. Animal. 2007;1(1):21–28.

36. Westell R, Quaas R, Van Vleck LD. Genetic groups in an animal model. Journal of dairy science. 1988;71(5):1310–1318.

37. Quaas RL. Additive genetic model with groups and relationships. Journal of Dairy Science. 1988;71(5):1338–1345.

38. Legarra A, Aguilar I. Tools to refine unknown parent group definitions. Journal of Dairy Science. 2025;108(2):1747–1756.

39. Farias FH, Johnson GS, Taylor JF, Giuliano E, Katz ML, Sanders DN, et al. An *ADAMTS17* splice donor site mutation in dogs with primary lens luxation. Investigative ophthalmology & visual science. 2010;51(9):4716–4721.

40. Downs L, Bell J, Freeman J, Hartley C, Hayward L, Mellersh C. Late-onset progressive retinal atrophy in the Gordon and Irish Setter breeds is associated with a frameshift mutation in *C2orf71* . Animal Genetics. 2013;44(2):169–177.

41. Bennett NE, Mirabal Torres SE, Gray PB. Exploratory content analysis of direct-to-consumer pet genomics: What is being marketed and what are consumers saying? Plos one. 2022;17(1):e0261694.

42. The Royal Kennel Club. Understanding and using your dog’s DNA test results; 2026. https://www.thekennelclub.org.uk/health-and-dog-care/health/getting-started-with-health-testing-and-screening/dna-testing/breeding-advice-for-dna-test-results/. Accessed 4 September 2026.

43. The Royal Kennel Club. Health Test Results Finder (Dog look up); 2026. https://www.royalkennelclub.com/search/health-test-results-finder/. Accessed 4 September 2026.

44. R Core Team. R: A Language and Environment for Statistical Computing. Vienna, Austria; 2024. Available from: https://www.R-project.org/.

45. Wickham H, Averick M, Bryan J, Chang W, McGowan LD, François R, et al. Welcome to the tidyverse. Journal of Open Source Software. 2019;4(43):1686.

46. Kinghorn BP. A simple method to detect a single gene that determines a categorical trait with incomplete penetrance. In: 50 Years of DNA: Proceedings of the Fifteenth Conference, Association for the Advancement of Animal Breeding and Genetics. Melbourne, Australia; 2003. p. 103–106. 7–11 July.

47. Kelley CT. Solving nonlinear equations with Newton’s method. SIAM; 2003.

48. Calus M, Bouwman A, Hickey J, Veerkamp R, Mulder H. Evaluation of measures of correctness of genotype imputation in the context of genomic prediction: a review of livestock applications. animal. 2014;8(11):1743–1753.

49. Lin P, Hartz SM, Zhang Z, Saccone SF, Wang J, Tischfield JA, et al. A new statistic to evaluate imputation reliability. PloS one. 2010;5(3):e9697.

50. Ramnarine S, Zhang J, Chen LS, Culverhouse R, Duan W, Hancock DB, et al. When does choice of accuracy measure alter imputation accuracy assessments? PloS one. 2015;10(10):e0137601.

51. van Rijsbergen. Information Retrieval. Butterworths; 1979.

52. Siblini W, Fréry J, He-Guelton L, Oblé F, Wang YQ. Master Your Metrics with Calibration. In: Advances in Intelligent Data Analysis XVIII. Springer International Publishing; 2020. p. 457–469.

53. Palmer C, Pe’er I. Bias characterization in probabilistic genotype data and improved signal detection with multiple imputation. PLoS genetics. 2016;12(6):e1006091.

54. MacCluer JW, VandeBerg JL, Read B, Ryder OA. Pedigree analysis by computer simulation. Zoo biology. 1986;5(2):147–160.

55. Kutzler MA, Mohammed HO, Lamb SV, Meyers-Wallen VN. Accuracy of canine parturition date prediction from the initial rise in preovulatory progesterone concentration. Theriogenology. 2003;60(6):1187–1196.

56. Uchańska O, Ochota M, Eberhardt M, Nizański W. Dead or alive? A review of perinatal factors that determine canine neonatal viability. Animals. 2022;12(11):1402.

57. Bates D, Mächler M, Bolker B, Walker S. Fitting linear mixed-effects models using lme4. Journal of statistical software. 2015;67:1–48.

58. Lewis T, Windig JJ. Dog breeds: towards genomic management of populations with a high incidence of genetic defects. In: Genomic management of animal genetic diversity. Wageningen Academic; 2017. p. 179–205.

59. The Kennel Club. Irish Setter breed health and conservation plan. The Kennel Club; 2016. https://irishsetterclubofscotland.co.uk/wp-content/uploads/2022/02/Irish-Setter_BHCP.pdf. Accessed 11 July 2026.

60. Wade CM, Nuttall R, Liu S. Comprehensive analysis of geographic and breed-purpose influences on genetic diversity and inherited disease risk in the Doberman dog breed. Canine Medicine and Genetics. 2023;10(1):7.

61. Garcia AM, Diaz-Papkovich A, Sillon G, D’Agostino D, Chong AL, Chong G, et al. Using the ancestral recombination graph to study the history of rare variants in founder populations. bioRxiv. 2025;p. 2025–03.

62. The Royal Kennel Club. Bull Terrier; 2026. https://www.royalkennelclub.com/search/breeds-a-to-z/breeds/terrier/bull-terrier/. Accessed 4 September 2026.

63. The Royal Kennel Club. Bull Terrier (Miniature); 2026. https://www.royalkennelclub.com/search/breeds-a-to-z/breeds/terrier/bull-terrier-miniature/. Accessed 4 September 2026.

64. Berry D, McParland S, Kearney J, Sargolzaei M, Mullen MP. Imputation of ungenotyped parental genotypes in dairy and beef cattle from progeny genotypes. Animal. 2014;8(6):895–903.

65. Berry D, McHugh N, Randles S, Wall E, McDermott K, Sargolzaei M, et al. Imputation of non-genotyped sheep from the genotypes of their mates and resulting progeny. animal. 2018;12(2):191–198.

66. Hollands GJ, French DP, Griffin SJ, Prevost AT, Sutton S, King S, et al. The impact of communicating genetic risks of disease on risk-reducing health behaviour: systematic review with meta-analysis. bmj. 2016;352.

67. Hayakawa S, Marian V. Communicating risk: How relevant and irrelevant probabilistic information influences risk perception in medical decision-making. Cognitive, Affective, & Behavioral Neuroscience. 2023;23(3):678–690.

68. Miller DT, Lee K, Gordon AS, Amendola LM, Adelman K, Bale SJ, et al. Recommendations for reporting of secondary findings in clinical exome and genome sequencing, 2021 update: a policy statement of the American College of Medical Genetics and Genomics (ACMG). Genetics in Medicine. 2021;23(8):1391–1398.

69. Skipper A. A new future for dog breeding. The Royal Kennel Club; 2026. https://www.royalkennelclub.com/media/cbsnewyj/rkc_report_2026_full-version.pdf. Accessed: 20 April 2026.

70. Cunningham L. The case against dog breed discrimination by homeowners’ insurance companies. Connecticut Insurance Law Journal. 2004;11.

71. Williams A, Williams B, Hansen CR, Coble KH. The impact of pet health insurance on dog owners’ spending for veterinary services. Animals. 2020;10(7):1162.

72. RSPCA, Scottish SPCA, Ulster SPCA. The Animal Kindness Index: Key findings; 2023. https://www.rspca.org.uk/whatwedo/latest/kindnessindex/2023/keyfindings. Accessed 4 September 2026.

73. The People’s Dispensary for Sick Animals. Pet Animal Wellbeing (PAW) Report. The People’s Dispensary for Sick Animals; 2023. https://www.pdsa.org.uk/media/13976/pdsa-paw-report-2023.pdf. Accessed 11 Oct 2026.

74. of the ‘Cost-of-Living Crisis’ on UK-Based Owners of Pandemic Puppies I. Brand, Claire L and Belshaw, Zoe and O’Neill, Dan G and Bryson, Gina T and Packer, Rowena MA. Animals. 2026;16(17):2656.

75. Hollinshead F, Ontiveros M, Burns J, Magee C, Hanlon D. Factors influencing parentage ratio in canine dual-sired litters. Theriogenology. 2020;158:24–30.

76. Legarra A, Christensen OF, Vitezica ZG, Aguilar I, Misztal I. Ancestral relationships using metafounders: finite ancestral populations and across population relationships. Genetics. 2015;200(2):455–468.

77. Macedo F, Astruc J, Meuwissen T, Legarra A. Removing data and using metafounders alleviates biases for all traits in Lacaune dairy sheep predictions. Journal of Dairy Science. 2022;105(3):2439–2452.

78. Moses L, Niemi S, Karlsson E. Pet genomics medicine runs wild. Nature. 2018;559(7715):470–472.

79. Gershony L, Oberbauer A, Clark LA. Review of the current state of genetic testing - a living resource. AKC Canine Health Foundation; 2020. https://dogwellnet.com/applications/core/interface/file/attachment.php?id=5247. Accessed 14 October 2026.

